# Dispersive currents from narrow windows of time explain patterns of population connectivity in an ecologically and economically important fish

**DOI:** 10.1101/2022.10.11.511760

**Authors:** Claire Schraidt, Amanda S. Ackiss, Wesley A. Larson, Mark D. Rowe, Tomas O. Höök, Mark R. Christie

**Affiliations:** Department of Forestry and Natural Resources, Purdue University; 715 W. State St., West Lafayette, Indiana 47907-2054 USA; Wisconsin Cooperative Fishery Research Unit, College of Natural Resources, University of Wisconsin-Stevens Point, 800 Reserve Street, Stevens Point, WI 54481, USA; U.S. Geological Survey, Great Lakes Science Center, 1451 Green Rd, Ann Arbor, MI 48105, USA; National Oceanographic and Atmospheric Administration, National Marine Fisheries Service, Alaska Fisheries Science Center, 17109 Point Lena Loop Road, Juneau, AK, 99801 USA; NOAA Great Lakes Environmental Research Laboratory, 4840 South State Rd., Ann Arbor MI 48108 USA; Illinois-Indiana Sea Grant, Purdue University, West Lafayette, IN 47907; Department of Biological Sciences, Purdue University; 915 W. State St., West Lafayette, Indiana 47907-2054 USA

**Keywords:** gene flow, larval dispersal, oceanographic currents, population connectivity, yellow perch

## Abstract

Identifying the drivers of population connectivity remains a fundamental question in ecology and evolution. Answering this question can be challenging in aquatic environments where dynamic lake and ocean currents, high variance in reproductive success, and above average rates of dispersal and gene flow can increase noise. We developed a novel, integrative approach that couples detailed biophysical models with eco-genetic individual-based models to generate ‘predictive’ values of genetic differentiation. We also used RAD-Seq to genotype 960 yellow perch (*Perca flavescens*), a species with an ∼30-day pelagic larval duration (PLD), collected from 20 sites circumscribing Lake Michigan. By comparing predictive and empirical values of genetic differentiation, we estimated the relative contributions for known drivers of population connectivity (*e.g.*, currents, behavior, PLD). For the main basin populations (*i.e.*, the largest contiguous portion of the lake), we found that high gene flow led to low overall levels of genetic differentiation among populations (*F_ST_* = 0.003). By far the best predictors of genetic differentiation were connectivity matrices that *1.* came from a specific week and year, and *2.* resulted in high population connectivity. Thus, these narrow windows of time during which highly dispersive currents occur are driving the patterns of population connectivity in this system. We also found that populations from the northern and southern main basin are slightly divergent from one another, while those from Green Bay and the main basin are highly divergent (*F_ST_* = 0.11). By integrating biophysical and eco-genetic models with genome-wide data, we illustrate that the drivers of population connectivity can be identified in high gene flow systems.

## Introduction

Identifying the spatial and temporal boundaries of freshwater and marine populations is critical for effective fisheries management and conservation (Begg et al., 1999; Carvalho & Hauser, 1995; Hixon et al., 2002). However, the delineation of aquatic populations can be challenging because many species are difficult to observe directly in their aquatic environments (Hedgecock et al., 2007). Furthermore, many fish and invertebrate populations are often connected by dispersal that occurs during a relatively cryptic pelagic larval stage throughout which most larvae are minuscule (∼1-5mm) and are nearly transparent, making them difficult to observe directly. This pelagic larval stage is ubiquitous; many freshwater fishes that inhabit large lakes and over 95% of all marine fishes have a pelagic larval stage as part of their life histories (Nelson et al., 2016). Being pelagic and with limited swimming ability, larvae can be transported on currents to locations that are hundreds of kilometers away from where they were spawned (Christie et al., 2010; Cowen et al., 2006; Williamson et al. 2016). On the other hand, behavioral adaptations, homing mechanisms, and a complex interplay of biophysical processes (including currents) can result in individuals returning to the same site from where they were spawned (Almany et al., 2007; Christie et al., 2010; D’Aloia et al., 2015). Thus, identifying the role that currents play in connecting aquatic populations remains central to the effective conservation and management of aquatic ecosystems (Burgess et al., 2014; Liggins et al., 2019).

Both theoretical and empirical studies have demonstrated the importance of currents in defining population connectivity in aquatic systems (Cowen et al., 2007; Cowen & Sponaugle, 2009; Pineda et al., 2007; Treml et al., 2008; Selkoe et al., 2010). A smaller, but still substantial, number of studies have identified relationships between oceanic currents (including biophysical models parameterized by current data) and genetic differentiation (Galindo et al., 2006; Krueck et al., 2020; Selkoe et al., 2016; Timm et al., 2020; White et al., 2010; Xuereb et al., 2018). However, most of these studies have relied on current patterns from single points in time or that were averaged across weeks, months, seasons, and years (but see Krueck et al., 2020) and few studies have identified the relationship between currents from shorter, specific time periods (*e.g.*, specific weeks) and population connectivity. Identifying whether currents from specific time periods play a large role in defining population connectivity is critical for: 1.) determining which currents to use for parameterizing connectivity matrices for use in theoretical or demographic models of population connectivity, 2.) understanding the ecological (*e.g.*, dispersal) and evolutionary (*e.g.*, gene flow) linkages among populations in space and time, and 3.) understanding patterns of genetic differentiation in aquatic systems. With respect to this last point, if currents from specific, and ostensibly narrow, windows of time play a large role in determining patterns of genetic differentiation in aquatic systems, then this result could also help explain the ‘chaotic genetic patchiness’ often described in marine systems. Chaotic genetic patchiness is commonly defined as unexpected patterns of genetic differentiation that are observed over small spatial scales and are not stable in time (*sensu* Broquet et al., 2013; Johnson & Black, 1982). Diverse and multifaceted drivers of chaotic genetic patchiness have been proposed, including genetic drift, high variance in reproductive success (a type of drift), and kinship (Iacchei et al., 2013). However, one relatively simple explanation for these patterns is that currents that connect local populations are highly dynamic and that only currents from certain time periods are responsible for most of the dispersal among populations in a given year.

There are several challenges with identifying specific currents from narrow windows of time as the drivers of population connectivity. First, high-resolution oceanographic data and coupled biophysical models are required from numerous time points. Thus, high-quality oceanographic data are prerequisite. Second, genetic differentiation is often used as a proxy for population connectivity, yet contemporary evolutionary processes (*e.g.*, genetic drift) and past evolutionary legacies (*e.g.*, signals of bygone selection) can confound genetic estimates of population connectivity (Waples, 1998; Waples & Gaggiotti, 2006; Whitlock & McCauley, 1999). Third, there is no standardized approach for correlating genetic estimates of population connectivity with oceanographic estimates of population connectivity (but see Krueck et al., 2020; White et al., 2010). One potential solution to this last challenge is to create species and system-specific eco-genetic individual-based models (Dunlop et al., 2009) that can use connectivity matrices derived from oceanographic biophysical models as input and return pair-wise genetic distance matrices as output (*e.g.,* Krueck et al., 2020). With this approach, empirically derived estimates of genetic differentiation (*e.g.*, *F_ST_*) obtained from genotyping individuals from multiple sites can be compared to eco-genetic model-derived estimates of genetic differentiation derived from spatially and temporally relevant oceanographic data. Here, we use this integrative approach across a 500 km latitudinal gradient to examine patterns of population connectivity in an ecologically and economically important fish species found throughout the Laurentian Great Lakes (hereafter Great Lakes).

In many ways, the Great Lakes have abiotic and biotic conditions that mirror temperate marine environments. High variability in near-shore currents, water temperature, and juvenile recruitment (Pritt et al., 2014) are just some of the characteristics shared between these systems. Perhaps most importantly, many Great Lakes fishes and invertebrates have a pelagic larval duration on the order of 30 to 40 days, high fecundity, high larval and juvenile mortality rates, and the potential for large population sizes – all characteristics shared with many marine species (Brazo et al., 1975; Forney, 1971; Ludsin et al., 2014; Pritt et al., 2014). Thus, insights gained from studies in the Great Lakes may be applicable to many marine systems and vice versa. In this study, we focused on yellow perch (*Perca flavescens*), an ecologically and economically important fish with an ∼ 30-day pelagic larval duration (Dettmers et al., 2005; Whiteside et al., 1985), collected from Lake Michigan. Lake Michigan spans a latitudinal gradient of nearly 500 km from north to south and has two large embayments, Green Bay and Grand Traverse Bay. While circulation patterns may vary substantially on an inter-annual basis, some general patterns are manifest. For example, during summer months, cyclonic gyres often form in the southern portion of the basin and along-shore currents from south to north dominate along the nearshore eastern portion of the lake (Beletsky et al., 2007; Höök et al., 2006). In the middle portion of the lake, the current patterns are often complex, but sometimes produce anticyclonic gyres that could act as barriers between southern and northern locations. Because substantial numbers of yellow perch larvae have been found in the middle of the lake (Dettmers et al., 2005), it is possible that these gyres could connect locations on the eastern and western sides of the lake. In the northern portion of the lake, the currents typically flow at much slower speeds than in the southern portion of the lake and the complex bathymetry and resulting embayments may support comparatively isolated aggregations of yellow perch.

Previous genetics work on yellow perch has revealed that throughout the Great Lakes, yellow perch have a shared evolutionary history (*i.e.*, the Great Lakes were most likely only colonized once). In fact, most yellow perch populations throughout the Great Lakes are dominated by a single mtDNA haplotype (Sepulveda-Villet et al., 2009). However, nuclear loci illustrate that there is a clear separation of yellow perch populations among each of the Great Lakes (Sepulveda-Villet & Stepien, 2012). Furthermore, the population connectivity of yellow perch within Lake Erie, which has been more extensively studied, demonstrate fine scale population genetic structure (Fraker et al., 2015; Sepulveda-Villet & Stepien, 2011), and there is no *a priori* reason to suspect that the patterns of population connectivity would be any less complex in Lake Michigan. One canonical study used a handful of nuclear loci to characterize the population structure of yellow perch from 5 sites in Lake Michigan (Miller, 2003) and found that sites in southern Lake Michigan appear to be genetically similar to one another and that there are substantial genetic differences between Green Bay and southern Lake Michigan. Nevertheless, fully characterizing the genetic structure of Lake Michigan yellow perch, as well as identifying the drivers of population connectivity in this system, has important implications for the successful conservation and management of species in this region and beyond (*e.g.*, stock identification, delineation of management units, identifying predictors of recruitment).

In this study, we sampled yellow perch from 20 sites circumscribing Lake Michigan. For every individual, we obtained genotypes at 9,302 SNP loci distributed throughout the yellow perch genome (Feron et al., 2020) using restriction site-associated DNA sequencing (RAD-Seq). We also used a Lagrangian particle tracking biophysical model to obtain high-quality current-derived connectivity matrices from 36 release dates spanning 6 years. With an integrated biophysical, eco-genetic model we varied particle release dates, pelagic larval duration, upward vertical swimming speed, and the number of generations of gene flow. Using these data, we asked three questions: 1.) What is the population structure of yellow perch throughout the main basin of Lake Michigan and Green Bay? 2.) How much does variation in the pelagic larval duration, vertical behavior of larvae, local population sizes, and number of generations of gene flow explain the empirically-derived estimates of genetic differentiation?, and 3.) How much do the specific weeks, months, and years of dispersal (*i.e.*, the release date used in the biophysical model) explain the empirically derived estimates of genetic differentiation? We find that there is significant population structure between Green Bay and the main basin of Lake Michigan and that population connectivity within the main basin is best explained by highly-dispersive currents from narrow windows of time.

## Materials and Methods

### Study Species and Sample Collection

Yellow perch remain one of the most ecologically and economically important species throughout the Great Lakes. They are an abundant near-shore fish species, serve as important predators of small fishes and invertebrates, and are themselves an important prey for larger fishes (Evans, 1986). Historically, yellow perch supported commercially-important fisheries throughout the Great Lakes region; in Lake Michigan alone, peak annual commercial harvest would now represent close to $16 million (US$) in dockside value and much more at retail. However, Lake Michigan yellow perch populations began declining during the late 1980’s and early 1990’s which led to closures of most yellow perch commercial fisheries. Relevant life history characteristics for yellow perch populations include: a pelagic larval duration on the order of 30 to 40 days (Dettmers et al., 2005; Whiteside et al., 1985), high fecundity (∼ 10,000 to 150,000 eggs/female), and type III survivorship (*i.e.*, high mortality during early life stages; Brazo et al., 1975; Forney, 1971). Yellow perch typically spawn during late spring to early summer when currents within Lake Michigan are often at their weakest (Beletsky et al., 1999). Adult yellow perch have modest home ranges and mark-recapture studies have demonstrated that most adult yellow perch and their congener, Eurasian perch (*Perca fluviatilis*), have high site fidelity, particularly with respect to spawning grounds (Bergek & Björklund, 2009; Böhling & Lehtonen, 1984; Glover et al., 2008; Schneeberger, 2000). Thus, the predominant form of population connectivity for yellow perch occurs during the pelagic larval stage. Similar to many marine fishes, yellow perch larvae may have some control over their dispersal trajectories simply by varying their vertical position within the water column (Graeb et al., 2004; Leis, 2006). As the larvae develop, they may also become better at swimming, such that a combination of active and passive dispersal mechanisms may ultimately dictate where individual yellow perch are located when they transition to a demersal life stage. Because yellow perch have many ecological and life-history characteristics in common with marine fishes, other Great Lakes fishes, and other commercially-important fishes (Ludsin et al., 2014; Pritt et al., 2014), they represent an ideal model system for studying patterns of population connectivity in aquatic systems.

Yellow perch were collected from 20 sites circumscribing Lake Michigan and Green Bay in 2018 and 2019 (Figure 1a, Table 1). In 2018, adults, (*i.e.*, individuals >100mm in total length; see Table S1 for size data) were sampled using 12 hour overnight multi-mesh gill net sets. Sampling during the 2018 season began in southern Lake Michigan in early March and continued through the end of July at the northernmost sites to account for variation in regional spawning times. Young-of-year fish (*i.e.*, juveniles) were sampled in September using beach seines. Individuals were classified as young-of-year based on size (<100mm total length). Sampling in both 2018 and 2019 was supplemented with assistance from agency partnerships, specifically the Michigan Department of Natural Resources, the Indiana Department of Natural Resources, the Wisconsin Department of Natural Resources, the Illinois Natural History Survey, and the Grand Traverse Band of Ottawa and Chippewa Indians, where sampling methods consisted of bottom trawls, gill netting, and creel surveys. Small portions of fin tissue (∼ 2cm x 2cm) were collected from every individual and stored in 95% non-denatured ethanol. A total of 1,376 individuals were sampled over two years from which a subset of 960 samples that maximized representation among collection sites were selected for genotyping (Table 1). Tissues were stored at −20°C upon arrival at Purdue University and held until DNA extraction.

**Figure 1:**
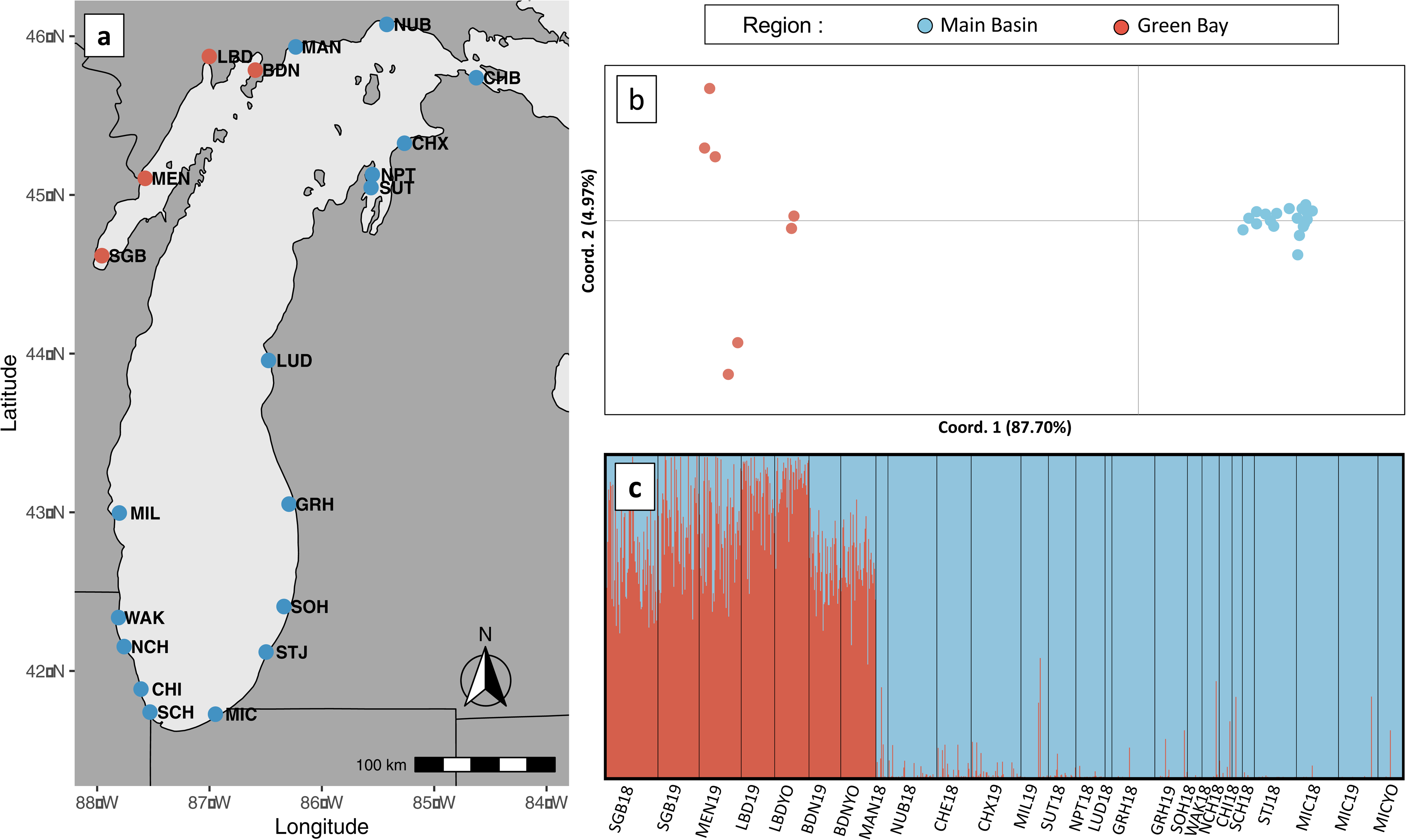
Sample collection sites and regional patterns of genetic differentiation. (a) A total of 960 yellow perch were collected and genotyped from 20 sites circumscribing Lake Michigan. (b) Principal coordinate analysis of pairwise *F_ST_* values illustrate substantial genetic differences between Green Bay and main basin yellow perch, where the first axis explains more than 87% of the total variation. (c) Results from STRUCTURE further illustrate the large genetic differences between Green Bay and main basin yellow perch. Notice that the proportion of main-basin ancestry increases for the Green Bay population closest to the main basin (BDN).

**Table 1:** Sampling details including collection site names, latitude, longitude, site name abbreviations (“Site ID”), region, year collected, age collected (yoy = “young of year”), and the number of individuals genotyped. A single individual from Algoma, Wisconsin was also genotyped but not used in any subsequent analyses (not shown).

|  |  |  |  |  | Year | Age | Number |
| --- | --- | --- | --- | --- | --- | --- | --- |
| Site Name | Latitude | Longitude | Site ID | Region | Collected | Collected | Sequenced |
| Michigan City | 41.72750 | -86.94735 | MICYO | Main Basin | 2018 | yoy | 28 |
|  |  |  | MIC19 | Main Basin | 2019 | adult | 46 |
|  |  |  | MIC18 | Main Basin | 2018 | adult | 50 |
| Saint Joseph | 42.11914 | -86.49644 | STJ18 | Main Basin | 2018 | adult | 50 |
| South Haven | 42.40596 | -86.33752 | SOH18 | Main Basin | 2018 | adult | 18 |
| South Chicago | 41.74067 | -87.53000 | SCH18 | Main Basin | 2018 | adult | 16 |
| Chicago | 41.88567 | -87.61000 | CHI18 | Main Basin | 2018 | adult | 13 |
| North Chicago | 42.15350 | -87.76000 | NCH18 | Main Basin | 2018 | adult | 15 |
| Waukegan | 42.38700 | -87.78000 | WAK18 | Main Basin | 2018 | adult | 20 |
| Grand Haven | 43.05141 | -86.29291 | GRH19 | Main Basin | 2019 | adult | 38 |
|  |  |  | GRH18 | Main Basin | 2018 | adult | 50 |
| Ludington | 43.95713 | -86.47504 | LUD18 | Main Basin | 2018 | adult | 10 |
| Milwaukee | 42.99538 | -87.80253 | MIL19 | Main Basin | 2019 | adult | 32 |
| Suttons Bay | 45.04650 | -85.56199 | SUT18 | Main Basin | 2018 | adult | 32 |
| Northport | 45.12820 | -85.55028 | NPT18 | Main Basin | 2018 | adult | 34 |
| Charlevoix | 45.32571 | -85.26504 | CHX19 | Main Basin | 2019 | adult | 59 |
| Cheboygan | 45.73871 | -84.62552 | CHB18 | Main Basin | 2018 | adult | 40 |
| Naubinway | 46.07465 | -85.42233 | NUB18 | Main Basin | 2018 | adult | 72 |
| Manistique | 45.93287 | -86.23245 | MAN18 | Main Basin | 2018 | adult | 15 |
| Big Bay de Noc | 45.78659 | -86.59284 | BDNYO | Green Bay | 2019 | yoy | 41 |
|  |  |  | BDN19 | Green Bay | 2019 | adult | 40 |
| Little Bay de Noc | 45.87262 | -87.00212 | LBDYO | Green Bay | 2019 | yoy | 41 |
|  |  |  | LBD19 | Green Bay | 2019 | adult | 40 |
| Menominee | 45.10461 | -87.57281 | MEN19 | Green Bay | 2019 | adult | 49 |
| South Green Bay | 44.61706 | -87.95805 | SGB19 | Green Bay | 2019 | adult | 50 |
|  |  |  | SGB18 | Green Bay | 2018 | adult | 60 |
|  |  |  |  |  |  | Total = | 959 |

### Molecular Methods

DNA was isolated from fin tissue with Qiagen DNeasy® Blood & Tissue Kits. Following an overnight (∼14 hour) tissue digestion with Proteinase K incubated at 56°C, DNA was extracted in plates using standard kit protocols and eluted from the silica membrane with 200uL Tris Low-EDTA buffer. Extractions were quantified using a Quant-it™ PicoGreen® dsDNA Assay (Invitrogen, Waltham, MA), and DNA was normalized to a quantity of 200ng or approximately 20ng/µL.

Libraries for restriction site-associated DNA (RAD) sequencing were prepared following the BestRAD protocol (Ali et al., 2016). Normalized DNA was digested with the restriction enzyme *SbfI* followed by ligation with indexed adaptors. Barcoded libraries were pooled into master libraries of 96 individuals and fragmented to ∼300-500bp with 12 30s cycles in a Q500 sonicator (Qsonica, Newtown, CT). Fragmented DNA was bound to Dynabeads™ M-280 Streptavidin magnetic beads (Invitrogen) and washed with buffer to remove non-target fragments. Following purification with AMPure XP beads (Beckman Coulter, Brea, CA), master libraries were passed in series through the NEBNext® Ultra™ DNA Library Prep Kit for Illumina® at the End Prep step for 1.) end repair and ligation of master library barcodes, 2.) a 250-bp insert size-selection, and 3.) a 12-cycle PCR enrichment. Successful size-selection and enrichment were confirmed with visualization of products on a 2% agarose E-Gel (Invitrogen). Products underwent a final AMPure XP purification clean-up followed by quantification with a Qubit® 2.0 Fluorometer. A total of 10 master libraries, each containing 96 individually barcoded samples, were sent to Novogene (Sacramento, CA) for PE150 sequencing on one lane of the Illumina Novaseq S4 platform.

Raw Illumina RAD sequence reads were processed using the STACKS v2.54 (Rochette et al., 2019) software pipeline. Reads were cleaned and demultiplexed by barcode using the STACKS subprogram process_radtags. Sequences were demultiplexed by barcode, filtered for Illumina quality score and enzyme cut-site, and trimmed to 140 base pairs to reduce tail-end sequencing errors (parameter flags = --filter_illumina, --bestrad, -t 140). The resulting filtered, individually-assigned reads were aligned to the yellow perch reference genome (*P. flavescens* PFLA_1.0 assembly, GenBank accession GCA_004354835.1; Feron et al., 2020) with bowtie2 (Langmead & Salzberg, 2012; Langmead et al., 2019) (parameter flag = --very-sensitive). Single nucleotide polymorphisms (SNPs) were called from reference aligned paired end reads with the STACKS subprogram gstacks (parameter flag = --rm-unpaired reads) and individuals were genotyped at each identified SNP. The gstacks output files, which contain consensus sequences at each identified locus, as well as individual genotype data, were filtered through the STACKS subprogram populations. SNPs that were genotyped in less than 30% of individuals were discarded, and analysis was limited to the first SNP per locus (parameter flag = -r 0.3) and results were exported in variant call format (VCF). SNPs were then filtered using vcftools v0.1.9 (Danecek et al., 2011). Post-STACKS filtering largely followed the same workflow published in Gehri et al., 2021. Briefly, filtering consisted of 1.) removing SNPs that were genotyped in fewer than 70% of individuals (Figure S1), (2) filtering out individuals with > 70% missing loci, and (3) removing loci with a minor allele frequency (maf) of less than 0.01. We next used HDPlot (McKinney et al., 2017) to remove loci with a read ratio deviation greater than 5 and less than −5. A custom python script was then used to select SNPs with the highest allele frequency at each position, and, in a final filtering step, SNPs with a maf less than 0.05 were removed. The resulting vcf file was converted to GENEPOP and STRUCTURE format using PGDSpider (Lischer & Excoffier, 2012).

### Population genomics

Collection site summary statistics including observed heterozygosity (*H_O_*) and expected heterozygosity (*H_E_*) were calculated in R 4.0.2 (R Core Team, 2021) using the package adegenet v.2.1.1 (Jombart, 2008). Allelic richness (*Ar*) was calculated using the R package hierfstat v0.5.7 (Goudet, 2005). We used the R package HardyWeinberg (Graffelman, 2015) to test for deviations from Hardy Weinberg Equilibrium using exact tests on each locus within each collection site. A Bonferroni correction based on the number of polymorphic loci within each collection site sample was used to identify loci out of equilibrium. A permutation test was used to determine whether regional genetic diversity estimates (*H_O_* and *Ar*) between Green Bay and the main basin differed from one another (see SI text and Figure S2 for details).

Cluster analyses were performed on all sites as well as separately within the main basin and Green Bay populations. The Bayesian clustering method implemented in STRUCTURE v2.3.4 (Pritchard et al., 2000) was first used to determine population structure present among all sampled locations. For STRUCTURE analyses with all samples, the optimal inferred cluster (K) was determined using the delta K method (Evanno et al., 2005). Runs consisted of an initial burn in period of 50,000 Markov Chain Monte Carlo (MCMC) iterations followed by 50,000 iterations for each inferred cluster. Analyses were performed with K = 1-30 clusters and replicated five times for each value of K. For separate main basin and Green Bay STRUCTURE analyses, we employed admixture and correlated allele frequency models, as this approach is most appropriate when subtle population structure is expected (Falush et al., 2003; Hubisz et al., 2009). Analyses were performed for K=1-20 for the 19 main basin sites and for K=1-8 for the 7 Green Bay sites, representing the total number of collection sites plus one, and replicated five times for each inferred cluster. As above, all runs consisted of an initial burn in period of 50,000 Markov Chain Monte Carlo (MCMC) iterations followed by 50,000 iterations for each inferred cluster.

Pairwise *F_ST_* between populations and 95% confidence intervals (calculated via bootstrapping, *n* = 1000) were calculated in hierfstat v0.5.7 (Goudet, 2005). Pairwise *F_ST_* values were output as a genetic distance matrix and exported for analysis in GenAlEx v6.5 (Peakall & Smouse, 2006), where principal coordinate analysis (PCoA) was performed in order to visualize genetic differentiation among populations. In both the main basin and Green Bay populations, principal components analysis (PCA) was used to further support the results of both STRUCTURE and PCoA for both the Green Bay and main basin populations. Allele frequencies were scaled using the scalegen function in adegenet, and PCAs were run using these scaled matrices with the base R function prcomp. For the STRUCTURE and PCA analyses, we used the LD.thin function in the R package gaston 1.5.7 (Perdry & Dandine-Roulland, 2020) with a threshold of 0.1 and max.dist of 500,000 to remove loci that were in linkage disequilibrium. This procedure retained 5,807 loci that resolved population structure marginally better than the full data set. Using the Green Bay and main basin pairwise *F_ST_* values, we also created isolation-by-distance plots where distances were calculated as the nearest along-shore distance between collection sites. We used a mantel test in GenAlEx to test for a positive relationship between distance and *F_ST_*.

### Biophysical models

For the biophysical model, we used a Lagrangian particle tracking model previously developed to study the transport of larval cod (Churchill et al., 2011; Huret et al., 2007), where three-dimensional current velocities and turbulent diffusivity were output from the application of the Finite Volume Community Ocean Model (FVCOM). A random-walk scheme for spatially varying vertical diffusivity was used, including a vertical floating/sinking/swimming velocity (Gräwe, 2011; Rowe et al., 2016). Particles were designated to be either 1.) neutrally buoyant or 2.) have an upward vertical swimming velocity of 0.0003 m/s. The Lagrangian particle tracking simulations were forced by output from FVCOM simulation of Lake Michigan-Huron (Anderson & Schwab, 2013) incorporating exchange currents in the Straits of Mackinac. Horizontal grid resolution varied with finer resolution nearshore and in regions with complex coastlines (*e.g.*, 100 m in the Straits of Mackinac to 2.5 km in the center of the lakes), and each horizontal grid was discretized into 20 terrain-following (sigma) layers. Additional details can be found in Supporting Information.

To generate connectivity matrices, the probability of transport from region *i* to region *f* was calculated as *N_if_* /*N_i_*, where *N_if_* is the number of particles initiated in region *i* that were within region *f* at the end of the simulation, and *N_i_* is the total number of particles that were initiated in region *i* (Figure S3). Based on the FVCOM and particle tracking models, connectivity matrices were developed for six years: 2014-2019. The sensitivity of the connectivity matrices to model assumptions were evaluated by considering scenarios of, 1.) vertical swimming behavior, and 2.) horizontal diffusion. Simple behavior scenarios were tested, including passive particle, upward swimming, and downward swimming. Scenarios with vertical swimming velocity were implemented by applying a deterministic vertical velocity in the vertical random walk turbulence scheme (Rowe et al., 2016), representing the combined effects of turbulence and directed swimming. Particles were initiated at the nodes of the unstructured FVCOM grid, at locations < 10 m deep (number of nodes = 2,246), consistent with nearshore spawning of yellow perch. Horizontal resolution was 200-600 m in nearshore areas where particles were initiated with 100 particles per node, uniformly distributed vertically through the water column. In scenarios with vertical swimming velocity set to zero, particles primarily remained distributed through the epilimnion, but some particles dispersed into the metalimnion in longer simulations. We also conducted scenarios with an upward swimming velocity sufficient to maintain particles within the epilimnion; an upward velocity of 0.0003 m/s was selected, which was considerably less than reported maximum horizontal swimming velocities of larval yellow perch of 0.03-0.046 m/s (reviewed by Höök *et al*. 2006), but sufficient to keep particles within the epilimnion (See SI text for details). We assigned a horizontal diffusion coefficient of 5.6 m^2^/s based on estimates from Lake Michigan (Thupaki et al., 2013). Models were run for three estimates of pelagic larval duration (30, 40, and 50 days; the mean and 10 days on either side; Beletsky et al., 2007), two upward swimming velocities (0 and 0.0003 m/s), 6 weekly release dates ranging from late May to early July (the peak estimated yellow perch spawning period; Starzynski & Lauer, 2015), and 6 years (2014-2019) resulting in a total of 216 biophysical model simulations (Table 2).

**Table 2:**
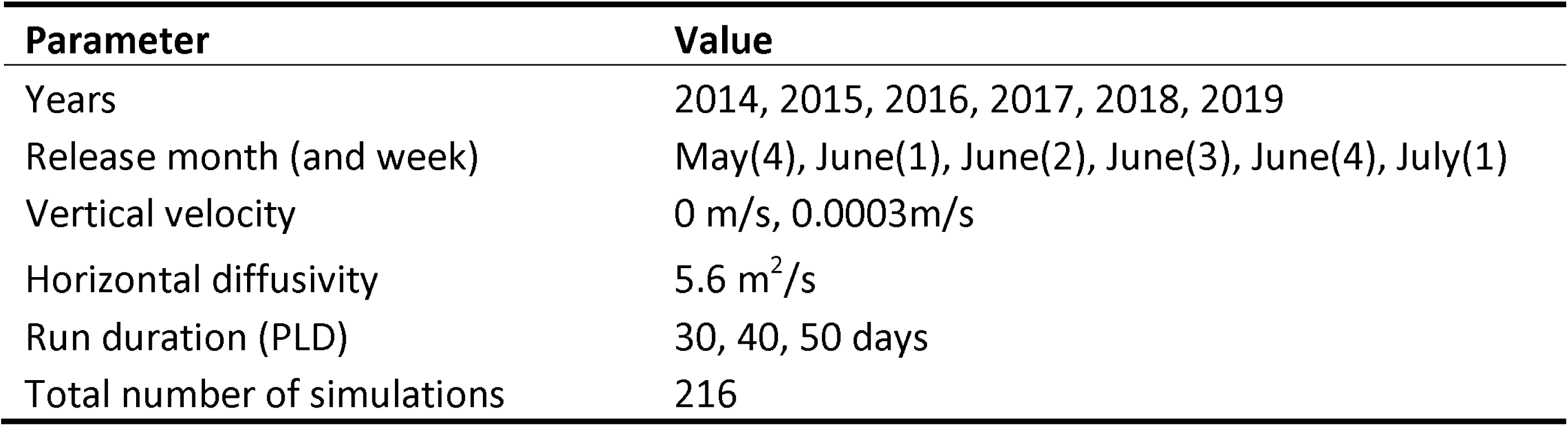
Parameter values used in the biophysical model for Lake Michigan yellow perch. A total of 216 different model runs were conducted where all possible combinations of 6 years, 6 particle release weeks (spanning 3 months), 2 vertical upward swimming velocities, and 3 run durations were examined. Run duration consisted of the time period from when the particles were first released to when the model was stopped and are thus analogous to varying the pelagic larval duration (PLD).

| Parameter | Value |
| --- | --- |
| Years | 2014, 2015, 2016, 2017, 2018, 2019 |
| Release month (and week) | May(4), June(1), June(2), June(3), June(4), July(1) |
| Vertical velocity | 0 m/s, 0.0003m/s |
| Horizontal diffusivity | 5.6 m<sup>2</sup>/s |
| Run duration (PLD) | 30, 40, 50 days |
| Total number of simulations | 216 |

### Eco-genetic models

To determine the extent to which currents could explain the empirically-estimated genome-wide levels of genetic differentiation, we created a spatially-explicit, forward-time individual-based model that simulates larval dispersal among populations and, importantly, is parameterized by the lake-wide biophysical connectivity matrices described above. The model was adapted from Christie et al. (2017) to incorporate our sample design (Figure 1) and yellow perch life history characteristics. The model was parameterized with 40 sites (hereafter: “local populations”) to mimic the grid design employed by the biophysical model (Figure S3). Each local population in the model was characterized by an average of 600-1,200 individuals, resulting in an average metapopulation size of 36,000 perch. Every individual was randomly assigned a sex (male or female) and was characterized by 100 independent (unlinked) single nucleotide polymorphisms (SNPs). At the beginning of each model run, all multi-locus genotypes were created in accordance with Hardy-Weinberg Equilibrium (HWE). After initializing populations, the model was characterized by the following steps: mortality, reproduction, larval dispersal, and recruitment. We assumed an average 20% mortality rate per year, following estimates for both yellow perch and for many coastal marine species (Wilberg et al., 2005). This process created age-structured populations with overlapping generations and a mean generation time of 4.8 years. During the mortality step, individuals were randomly removed from throughout the metapopulation without respect to local population. Within each local population, mortality rates were varied slightly each year using a random deviate from a normal distribution with a mean equal to the number of offspring needed for replacement and a standard deviation of 105. This process increased fluctuations in local population sizes and mimics population dynamics of Great Lakes perch populations (Irwin et al., 2009; Figure S4).

Because many aquatic organisms are characterized by high variance in reproductive success (Hedgecock & Pudovkin, 2011), we varied the number of offspring produced by each pair (with most pairs producing no offspring surviving to recruitment in a given year) using a gamma distribution with a shape parameter of 0.5 and a rate parameter of 0.1. Pairs were created by randomly pairing males and females within each local population without replacement. Offspring were created in strict accordance with Mendelian inheritance; at each locus, each offspring inherited one allele, chosen at random, from both parents. To simulate larval (*i.e.*, offspring) dispersal among the 40 local populations, we used connectivity matrices from the FVCOM biophysical model (see section above). A total of 216 connectivity matrices were available (Table 2). For each model run, we first selected a single connectivity matrix and applied the selected connectivity matrix to each local population in the model to determine the number of recruits originating from each of the 40 local populations. Specifically, we used a multinomial distribution specifying all 40 populations, the number of needed offspring for a local population to return to its local carrying capacity, and the connectivity matrix describing the probability of a recruit originating from each of *i* local populations. In practice, the multinomial distribution was implemented prior to reproduction so that we knew precisely how many offspring to create in each local population, which increased computational efficiency, however, the actual dispersal of individuals occurred after reproduction.

Because there is a fair amount of uncertainty associated with the demographic, life history, and dispersal characteristics of Lake Michigan yellow perch, we tested a total of 4,536 combinations of parameter values (Table 3), with each unique set of parameters replicated with 100 model runs. Thus, a grand total of 453,600 individual simulations were run on four high-performance computing nodes (256 cores, 1024 GB memory). Each set of parameters consisted of a combination of parameters specified by the biophysical or eco-genetic models (Table 3). Specifically, we selected one of three values for the number-of-years to run the eco-genetic model (50 years, 100 years, or 200 years), one of three pelagic larval durations (30 days, 40 days, or 50 days), one of four estimates of local population sizes (600, 800, 1000, 1200), one of two upward swimming speeds (0 and 0.0003 m/s), and for date-specific current data, one of 6 years (2014-2019), and one of 6 release weeks (the 4th week of May, the 1st through 4th weeks of June, and the 1st week of July) (Table 3), which correspond with known peaks of yellow perch spawning events in Lake Michigan (Starzynski & Lauer, 2015). We also wanted to test whether varying connectivity matrices within a single simulation would improve predictive ability. Thus, we also included model runs where the connectivity matrix was replaced for each year of the eco-genetic model. Specifically, we: 1.) randomly selected (with replacement) a connectivity matrix from the first 3 release weeks across all six years (2014-2019) for every year in the eco-genetic model, 2.) randomly selected a connectivity matrix from the last 3 release weeks across all six years (2014-2019) for every year in the eco-genetic model, and 3.) randomly selected a connectivity matrix from all 6 release weeks across all six years (2014-2019) for every year in the eco-genetic model (see sets 1-3 in Table 3). The first two scenarios were tested because the spawning window for yellow perch may be earlier (scenario 1) or later (scenario 2) than the entire 1.5-month window, but not well characterized by a single release year. The last scenario (scenario 3) tests whether averaging across all release weeks and years is a better predictor of connectivity than a single release week and year. Lastly, we performed a similar set of analyses where for each combination of parameters we randomly selected the release weeks (identical to scenarios 1-3), but kept the year fixed (*i.e.*, ran one year at a time). This procedure allowed us to test whether averaging over years provided better predictive value than connectivity matrices from a single, narrow point in time.

**Table 3:** Parameter values used in the eco-genetic individual-based model for Lake Michigan yellow perch, which was parameterized by connectivity matrices generated from a detailed biophysical model. Model years represents the number of years the eco-genetic model was run through all steps (*i.e.*, reproduction, dispersal, mortality). Matrix rows represent the individual release years, months, and weeks used for determining dispersal among local populations in the eco-genetic model. Each of the 216 connectivity matrices were tested separately. We also tested specific sets of connectivity matrices where connectivity matrices were allowed to vary each year within the eco-genetic model (*i.e.*, a connectivity matrix designated within each set was randomly selected with replacement each model year prior to the reproduction step). For each unique combination of input parameters (N=4,536) we ran 100 replicate simulations.

| Parameter | Value |
| --- | --- |
| Model years | 50, 100, 200 |
| Matrix years | 2014, 2015, 2016, 2017, 2018, 2019, 2014-2019 |
| Matrix release month (and week) | Individual: May(4), June(1), June(2), June(3), June(4), July(1) |
| Matrix release month (and week) sets | Set 1=[May(4), June(1), June(2)],<br>Set 2=[June(3), June(4), July (1)],<br>Set 3=[May(4) - July (1)] |
| Matrix vertical velocity | 0 m/s, 0.0003m/s |
| Matrix run duration (PLD) | 30, 40, 50 days |
| Local populations | 40 |
| Local population sizes | 600, 800, 1000, 1200 |
| Annual mortality rate | 0.2 |
| Variance in reproductive success | Gamma (alpha = 0.5, beta = 0.1) |
| Metapopulation carrying capacity | 40000 |
| Number of loci | 100 |
| Number of alleles per locus | 2 |
| Replicates | 100 |
| Total number of simulations | 453,600 |

At the end of each model run, we calculated pairwise genetic differentiation (unbiased *F_ST_*; Weir & Cockerham, 1984) among all pairs of populations. These pairwise estimates were derived entirely from the eco-genetic model (hereafter: “predictive values”) and were compared to the empirical *F_ST_* values derived from genotyping the field-collected samples (hereafter: “empirical values”). Thus, we obtained values of the exact same estimator calculated with two entirely independent approaches. Because the same estimator was used, a perfect fit between the predictive values and the empirical values would fall directly on a 1:1 line (*i.e.,* y = x). Thus, to estimate the accuracy and precision of the predictive values we used simple linear regression and calculated the slope and correlation (measured here as the coefficient of determination; adjusted *R^2^*) between the empirical and predictive values. To account for the possibility of non-linear relationships, we also used Spearman’s rank correlation coefficient, which returned qualitatively identical and nearly quantitatively identical results (Figure S5). To estimate the goodness of fit, we plotted the results of fitting a linear model to the predictive vs. empirical estimates for each set of parameters (averaged over 100 replicates) and plotted the mean correlation versus the mean slope across all unique combinations of parameter values. Parameter values resulting in high predictive power have correlation and slope values close to one and this approach allows us to isolate the effect of individual parameters against a background of thousands of different combinations of parameter values. For each parameter, we isolated the tested values (Table 3) that were in the top 20% of all simulations with respect to correlation and slope and calculated their relative contributions (*e.g.*, predictive values generated with connectivity matrices from 2016 resulted in many more predictions in the top 20% of all parameter values than those generated from 2014). Using more stringent criteria (*i.e.*, top 10% of all simulations) resulted in qualitatively similar results with more pronounced effects (*i.e.,* larger effects of release year and week). We also combined slope and correlation estimates into a single goodness of fit metric (see SI Methods) to examine the contributions of specific weeks and years and to examine the relationship between larval connectivity (measured as the number of grid cells with particles originating from cell *i*, averaged for all values of *i*), larval retention (measured as the number of particles that originated and remained in cell *i*, averaged for all values of *i*), and goodness of fit. Lastly, the percent of variation explained by each parameter was estimated as the correlation between a specific predictor and the goodness of fit between predictive and empirical *F_ST_*. The eco-genetic model and all downstream analyses were written in R version 4.0.2 (R Core Team, 2021).

## Results

All 960 individuals were sequenced, producing over 5 billion reads that resulted in an average of 5,463,100 paired-end reads per sample. Following filtering, 927 individuals from 26 collection sites (delineated by site, year, and young-of-year vs. adult) were genotyped at 9,302 loci (Table 1). Mean read depth of loci across individuals was 29x and mean missingness per individual was 7.6% (Figure S1). The estimates of global genetic diversity for heterozygosity and allelic richness were *H_o_* = 0.246, *H_e_* = 0.243, and *A_r_* = 1.248, respectively. Genetic diversity estimates were also calculated for each population (Table S2). An average of 36 loci (0.38%) were out of Hardy Weinberg equilibrium (HWE) within each population (range = 0.01 to 0.89%), while only a single locus was out of HWE across 70% or more of collection sites (>= 18/26 collection sites). Thus, we retained all loci for downstream analyses. Permutation tests revealed that genetic diversity (both observed heterozygosity and allelic richness) was statistically higher in Green Bay compared to main basin yellow perch (p-values < 0.001; Figure S2), however the absolute differences in heterozygosity and allelic richness (both < 0.06; Figure S2) were so small as to be of little biological significance.

Pairwise *F_ST_* values ranged from −0.001 to 0.148 (Table S3) where mean pairwise *F_ST_* was 0.003 among all main-basin populations, 0.018 among all Green Bay populations, and 0.11 between the main-basin and Green Bay populations. For STRUCTURE analysis across all collection sites, both mean likelihood values (L(K)) and ΔK suggested two optimal clusters (K=2) (Figure S6). This clustering reveals a distinct genetic split between Green Bay populations and main basin samples (Figure 1). The principal coordinates analysis for pairwise *F_ST_* values further supports this split, with main basin and Green Bay samples clustering out along the first coordinate, which explains 87.7% of the variance among the 26 collection sites (Figure 1). In Green Bay, STRUCTURE analysis revealed similar ΔK values for K = 2 and K = 3, however K = 3 appears to be better supported by the results from the principal coordinate and principal component analyses (Figure 2a,b,c, Figure S7). These three groupings represent Little Bay de Noc, Big Bay de Noc, and Southern Green Bay, respectively. When the main basin samples were run separately to determine fine-scale population structure, analysis revealed ΔK maxima at K=2, but visualizing these clusters revealed only subtle population structure between the northern and southern sites (Figure 2d,e,f; Figure S7).

**Figure 2:**
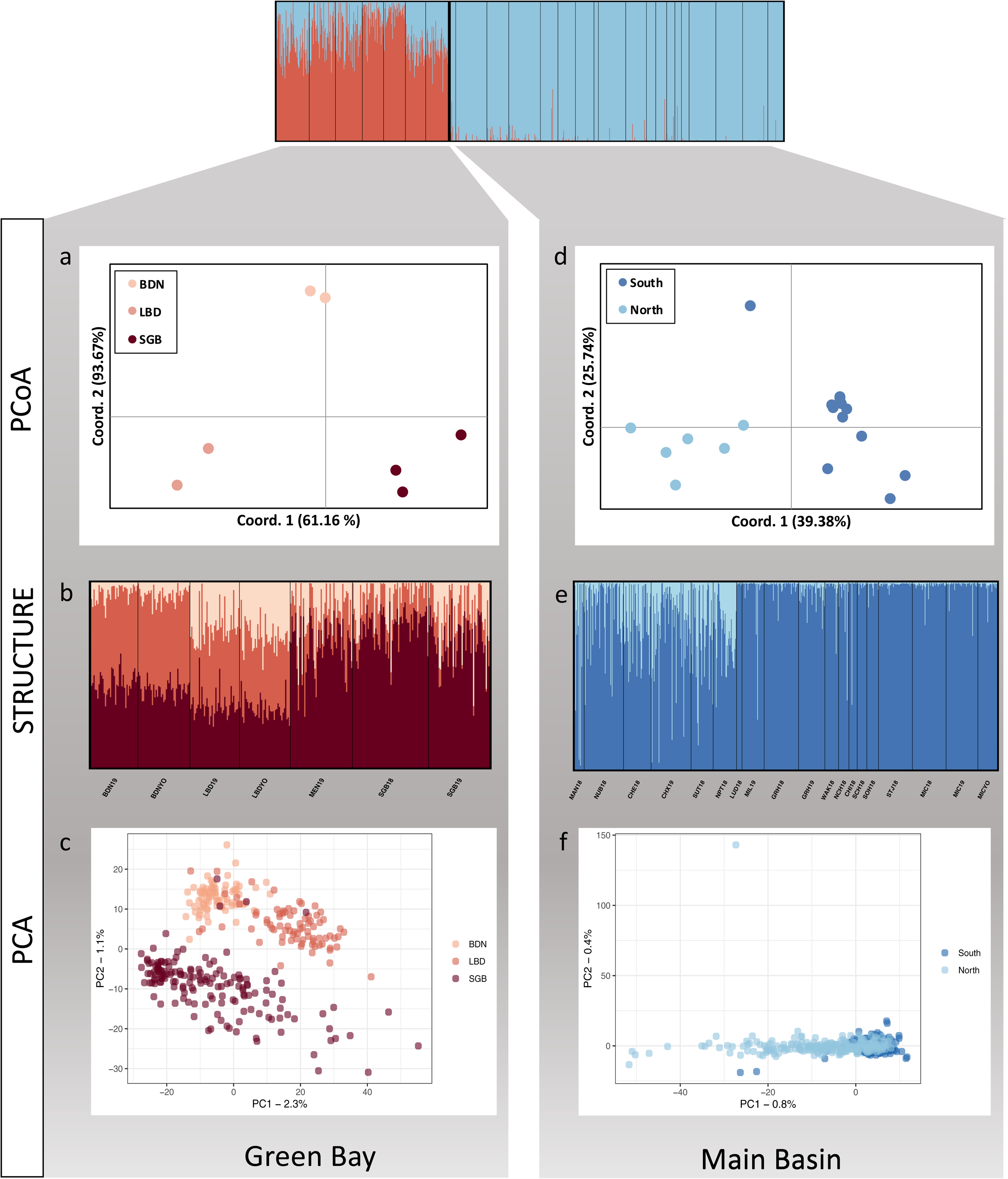
Patterns of genetic differentiation within each region. For Green Bay populations (red), principal coordinate analysis (PCoA) of pairwise *F_ST_* values (a), STRUCTURE output for K=3 (b), and principal component analyses (PCA) for individual genotypes (c) all reveal separation between Big Bay de Noc (BDN), Little Bay de Noc (LBD), and southern Green Bay (MEN and SGB). For the main basin populations, there is subtle population structure between northern and southern Lake Michigan collection sites as again determined by principal coordinate analysis of pairwise *F_ST_* values (d), STRUCTURE output for K=2 (e), and principal component analyses for all individual genotypes (f).

For Green Bay populations, there was a positive relationship between nearest along-shore distance and *F_ST_*(p = 0.05, *R^2^* = 0.29; Figure S8). For main basin populations, there was no relationship between nearest along-shore distance and *F_ST_* (p = 0.36, *R^2^* = 0.001; Figure S9) suggesting a need for additional metrics to explain patterns of genetic differentiation. By contrast, examining the relationship between the highest and lowest goodness of fit (*i.e.*, slope and correlations closest to one) between the current-derived, predictive *F_ST_* values and the sequence-derived, empirical *F_ST_* values, we found that the best predictors of empirical *F_ST_* occurred when there was high population connectivity for main basin populations (Figure 3a, b). Conversely, the worst fit occurred when there was high larval retention, and low population connectivity, resulting in model-based estimates of *F_ST_* that were twice as high as those found empirically (Figure 3c, d). This result was further validated when we examined the relationship between population connectivity (estimated directly from the connectivity matrices generated via the biophysical model) and the goodness of fit between predictive and empirical *F_ST_* (Figure 3e; slope = 2.10, *R^2^* = 0.31, p-value < 0.001). Stated differently, when lake-wide currents resulted in high among-population connectivity, those matrices more accurately predicted empirical *F_ST_*. We also saw that lower larval retention resulted in better predictive ability, though this pattern was not as strong as connectivity (Figure 3f; slope = −0.81, *R^2^*= 0.04, p-value < 0.001).

**Figure 3:**
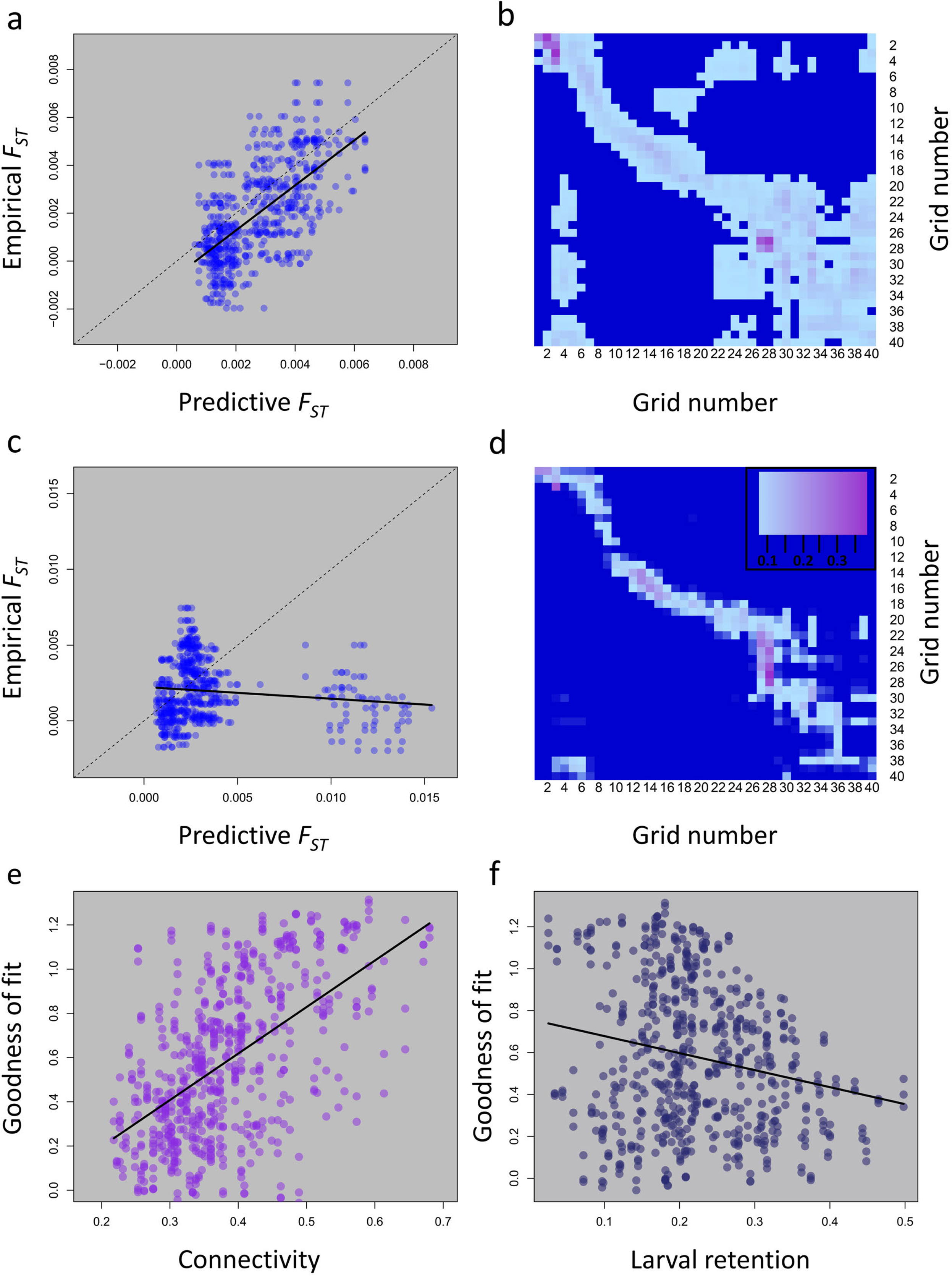
Relationships between predictive *F_ST_* values among main basin populations obtained from our eco-genetic model parameterized with biophysical current data and empirical estimates of *F_ST_* from 9,302 SNPs. The relationship between predictive *F_ST_* and empirical *F_ST_* for 10 replicated simulations for the connectivity matrix with the highest goodness of fit (a). A perfect fit (*i.e.,* perfect prediction) would result in all points laying directly on the 1:1 line (dashed line). The corresponding connectivity matrix with the best predictive ability (used to create predictive *F_ST_* values in panel a), occurred during a time period with high connectivity among neighboring sites (b). The relationship between predictive *F_ST_* and empirical *F_ST_* for 10 replicated simulations for the connectivity matrix with the lowest goodness of fit (c). Notice that many predictive pair-wise comparisons had *F_ST_* values 2-3 times higher than those observed empirically. The corresponding connectivity matrix, with low predictive ability, showed much lower connectivity among sites (d) (inset illustrates color scale used in b and d). In general, connectivity matrices with higher population connectivity had a higher predictive ability than matrices with low population connectivity (e). Similarly, connectivity matrices with higher rates of larval retention had a lower predictive ability, though this relationship was not as strong as for connectivity (f).

We next found that both the year and week that larvae were released were important drivers of predictive ability (Figure 4, Figure S10-S12). In particular, connectivity matrices from 2016 and the last week of June and first week of July were strong predictors of empirical estimates of *F_ST_*(Figure 4a, b) irrespective of other parameter values. Whether or not the connectivity matrices allowed for vertical, upward swimming also had a large effect (Figure 4c), where allowing for upward vertical swimming resulted in poorer predictions. In general, eco-genetic simulations parameterized with connectivity matrices from a 50-day PLD, versus a 30-day PLD had higher predictive value (Figure 4d). Varying the number of years that the eco-genetic model was run (50 vs. 100 vs. 200 years, Table 3), and thus the number of years of reproduction, mortality, and gene flow, had little effect on the goodness of fit between predictive and empirical values (Figure 4e). This result suggests that much of the present-day spatial genetic differentiation, at least as captured by RAD-Seq loci, is explained by processes that occur on contemporary time scales. Lastly, increasing the local population size for each of the 40 sites from 600 to 800 individuals improved predictive ability, but further increases in local population size had little effect (Figure 4e).

**Figure 4:**
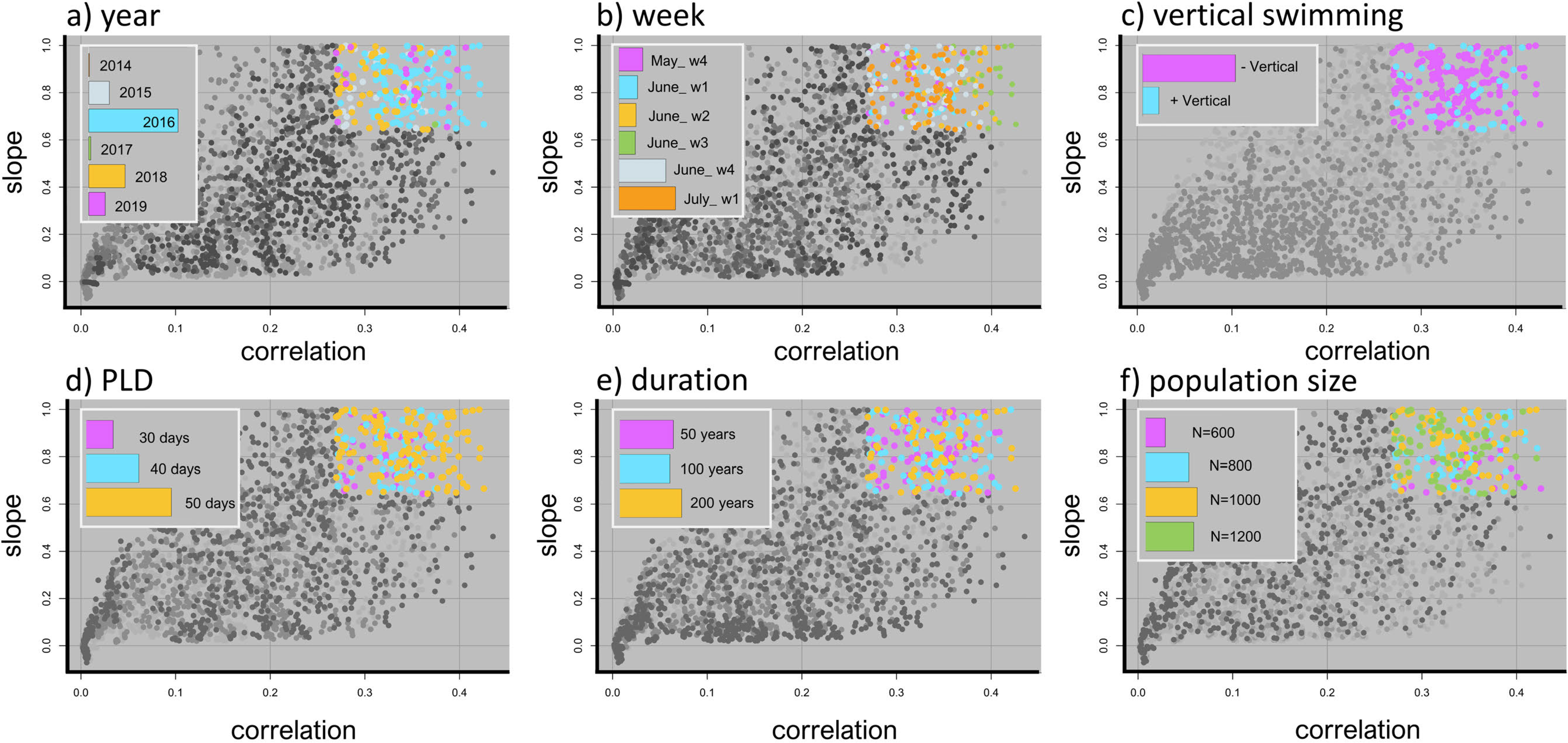
Drivers of population connectivity and genetic differentiation in main basin Lake Michigan yellow perch. Points represent the average correlation and slope values between predictive (generated from the integrated biophysical eco-genetic model) and empirical *F_ST_* for 100 simulations per unique set of parameter values (Table 3). A perfect fit would between predictive and empirical values would lie on the 1:1 line (y = x) and would have a correlation and slope equal to one. Colors represent the effect of particular parameters (across all sets of parameter values) for the top 20% of correlation and slope estimates (see Figure S10 for full color). Insets illustrate the relative contributions of particular parameter values contributing to the top 20% of model predictions. Across all parameters, the specific year and week that particles were released in the biophysical model had high predictive ability (a, b), as did vertical swimming ability (c). Pelagic larval duration, the number of years that the eco-genetic model was run (“duration”), and the local population size had lower predictive ability (d, e, f).

When we examined the joint effect of release year and week, we found that the specific week of release alone had the highest predictive ability of any single parameter (Figure 5). In general, the connectivity matrices from the 4^th^ week of June and the and 1^st^ week of July 2016 predicted the empirical estimates of genetic differentiation very well (Figure 5a, Figure S11). Conversely, other weeks in different years (*e.g.*, 2014) had very low predictive ability (Figure 5a). The observation that specific weeks and years had a much higher predictive ability than others suggests that narrow windows of time are important for determining patterns of population genetic connectivity. This result was further confirmed with the comparatively low predictive ability of eco-genetic output from models which were parameterized each year by randomly selected connectivity matrices across weeks and years (Figure S12). Thus, predictive *F_ST_* values from models where different connectivity matrices were used in each year of the eco-genetic simulation (*i.e.*, analogous to averaging currents across broader windows of time) were, on average, worse at predicting empirical estimates of genetic differentiation.

**Figure 5:**
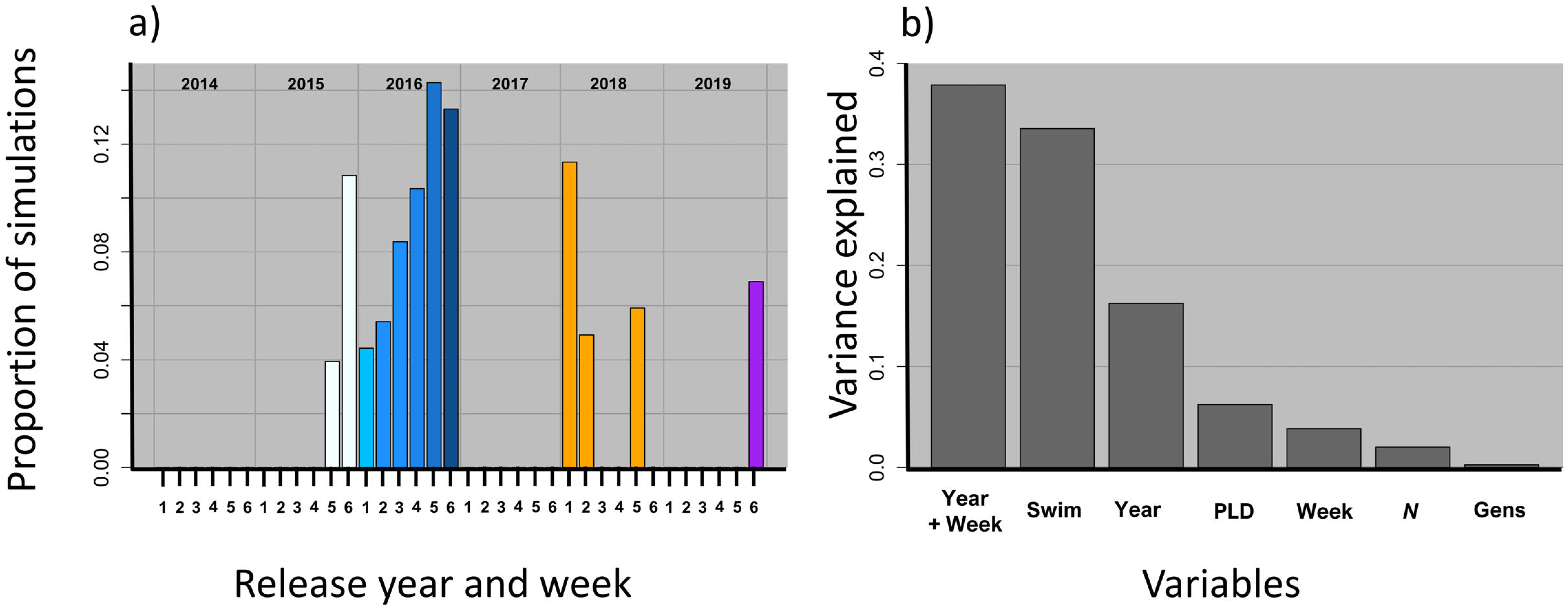
Relative contribution of dispersive currents from narrow windows of time and other drivers of genetic population connectivity in main basin yellow perch. A.) The effect of release year and week (across all sets of parameter values) for the top 20% of correlation and slope estimates illustrates that predictive values generated from connectivity matrices generated from specific weeks (especially those in 2016 and weeks 5 and 6) represented a higher proportion of all integrated biophysical eco-genetic simulations and thus have high predictive ability. The connectivity matrices with high predictive ability reflect time periods with highly dispersive currents (see Figure 3). Conversely, certain weeks had low predictive ability and never showed up in the top 20% of estimates (e.g., 2014). Release weeks correspond to: 1 = last week of May, 2 = 1^st^ week of June, 3 = 2^nd^ week of June, 4 = 3^rd^ week of June, 5 = 4^th^ week of June, 6 = 1^st^ week of July. B.) The percent of variance explained by various parameters (Table 3). The specific release year and week explained most of the variation, followed by vertical swimming behavior (Swim), year alone (Year), and pelagic larval duration (PLD). The week of release alone (Week), local population size (*N*), and number of generations that the eco-genetic model was run (Gens) explained the least amount of variation.

When we examined the percentage of variation explained by all parameters included in the eco-genetic model, we found that the particular week of reproduction and ensuing larval dispersal (*i.e.*, specific release date, year + week) explained the majority of differences in predictive ability (Figure 5b). Vertical swimming behavior, followed by year (*i.e.*, effect of year without respect to week) were the next best predictors (Figure 5b). The length of the pelagic larval duration, the local population size, and the number of years that the eco-genetic model was run for explained a much smaller percentage of the variation. Very long runs of the eco-genetic model (*i.e.*, 1000 or 2000 years) also did not have a large effect (Figure S13).

## Discussion

For an aquatic species with a pelagic larval stage, we demonstrated that genetic differentiation across a 500 km latitudinal gradient is best predicted by dispersive currents from narrow windows of time. Our integrative approach of combining oceanographic biophysical modeling with eco-genetic individual-based models and genomics illustrated that currents from particular weeks of particular years can be good predictors of genetic differentiation. Conversely, currents from other weeks can be very poor predictors of genetic differentiation and as little a difference in time as a single week can result in currents switching from being a strong predictor of genetic differentiation to a poor predictor and vice versa. Previous work has illustrated that ocean currents can be good predictors of genetic structure (Krueck et al., 2020; White et al., 2010), but, to our knowledge, no studies have illustrated that their predictive ability can vary over such a short time frame. Depending on the time frame and intervals over which oceanographic data are collected, this result means that previously published studies may have underestimated the role of currents in driving patterns of population connectivity in aquatic systems.

We further found that weeks that were good predictors of genetic differentiation were characterized by connectivity matrices with high levels of connectivity. Thus, in this system, the specific year or week (*e.g*., spring vs. summer) mattered less than whether that week (and the days of dispersal that followed afterwards) had highly dispersive currents. More generally, the observation that genetic differentiation in aquatic systems can be determined by dispersive currents from a specific period of time has broad implications. First, many examinations of population connectivity rely solely on data from biophysical models and rely on averaging or integrating current data over lengthy periods of time (e.g., Krueck et al., 2020; White et al., 2010). Here, we show that such analyses could be problematic because the realized population genetic connectivity may only occur during a narrow window of time. Thus, parameterizing connectivity matrices for use in theoretical or demographic models of population connectivity must be performed with caution. Second, the joint observations that 1.) specific weeks are accurate predictors of genetic differentiation, 2.) model runs employing multiple spawning dates (connectivity matrices from multiple weeks or years) performed worse than those from single release dates, and 3.) that the number of years for which the eco-genetic model was run had almost no effect on predictive ability (*e.g.*, Figure 4e, Figure S13) highlights the fact that, in this system, contemporary processes (and not evolutionary legacies) are likely to be driving patterns of genetic differentiation at most loci. Lastly, we speculate that in highly dynamic systems, such as many marine and freshwater environments, if most population connectivity occurs during a narrow window of time, then this result could explain, at least in part, many observations of chaotic genetic patchiness (Broquet et al., 2013; Johnson & Black, 1982).

Bolstering our interpretation that currents from specific weeks and years play a disproportionate role in determining patterns of population connectivity is the observation that 2015 and 2016 were particularly strong recruitment years for yellow perch in the main basin of Lake Michigan (Makauskas & Clapp, 2018). In turn, these year classes dominated the 2018 and 2019 perch population in the main basin (Makauskas & Clapp, 2018). Annual recruitment success of yellow perch in Lake Michigan and the entire Great Lakes region is positively related to spring-summer temperatures (Honsey et al., 2016). Of interest, 2016 (the year with best predictive ability) was particularly warm and 2014 (a year with very low predictive ability) was particularly cold. Annual differences in temperature also influence when yellow perch spawn and hatch. For example, Withers et al. (2015) documented earlier peak catches of recently hatched larval yellow perch in southern Lake Michigan during 2010, a particularly warm year, than during 2011, an intermediate thermal year. Such effects likely contribute to the observation that within years, particular weeks show differences in goodness of fit between empirical and predictive *F_ST_* values. Whether currents in particular years and weeks facilitate not only genetic population connectivity, but also recruitment, remains unknown as does the interaction between temperature, currents, and recruitment. This question could potentially be answered by an even sampling of different age classes, especially young-of-year, however we found it particularly challenging to collect younger age classes from the main basin due to their low abundance. Nevertheless, more work is needed to explore variation in yellow perch spawn timing, population connectivity, and subsequent recruitment.

Here, we have used an integrative approach to better understand the drivers of population connectivity in aquatic systems. This approach could be improved with higher-resolution oceanographic data, both in space and time. At least some of the unexplained variation between model-derived and empirical estimates of genetic differentiation could be due to a lack of resolution in the biophysical model, particularly with respect to near-shore currents (Gawarkiewicz et al., 2007). Furthermore, accurate estimates of local population sizes combined with accurate estimates of fecundity could improve both the biophysical model, in terms of more accurately dictating the number of particles to release at each site, and the eco-genetic model, in terms of accurately simulating gene flow and genetic drift. Releasing larger numbers of larvae per node and incorporating additional aspects of larval behavior such as horizontal swimming, particularly as yellow perch are strong swimmers towards the end of their PLD, could also help to improve the biophysical model (Kingsford et al. 2002; Leis & McCormick 2002). Additional data on spawning times and locations (*i.e.*, yellow perch may spawn in southern Lake Michigan earlier than yellow perch in northern Lake Michigan), may also bolster the ability to explain empirical patterns of genetic differentiation. Improving the eco-genetic model to incorporate site-specific demography and adult movement data, along with additional evolutionary forces such as mutation and selection, could also yield improvements. In short, any approaches that remove uncertainty in parameter estimates are likely to yield gains in prediction accuracy and aid in disentangling the relative contributions for drivers of population connectivity.

At a regional level, we found that Green Bay was highly genetically differentiated from the main basin (mean *F_ST_* = 0.11). This confirmation of previous work by Miller (2003) is most likely driven by the relatively low rates of water exchanged between the Green Bay and the main basin (Beletsky & Schwab, 2001) and further highlights the potential importance of managing Green Bay and main-basin yellow perch as distinct stocks. We also documented subtle population structure within the main basin itself (Figure 2) where southern and northern populations tend to cluster together. The genetic differences within the main basin, however, are much more comparable to marine systems, with mean pairwise *F_ST_* equal to 0.003. Surprisingly, yellow perch populations from Traverse Bay were much more similar to main-basin yellow perch populations than Green Bay yellow perch were to main basin populations. This result suggests that the local, and perhaps fine-scale, currents within Traverse Bay result in more connectivity with the main basin of Lake Michigan than Green Bay. From a population genetic perspective, it is interesting that Green Bay yellow perch show characteristics in common with fishes that lack a pelagic larval stage (higher *F_ST_*; isolation-by-distance; Figure S8), while main basin perch show patterns of genetic differentiation similar to many marine fishes (lower *F_ST_*; no pattern of isolation-by-distance Figure S9) (DeWoody & Avise, 2000; Martinez et al., 2018). Our three young-of-year samples each showed patterns of ancestry similar to their nearby adult samples (Figure 2, most easily seen in STRUCTURE plots) suggesting that high variance in reproductive success is not driving patterns of genetic differentiation in this system (*cf.* Christie et al., 2010), but more samples of fish across different age classes are needed.

In conclusion, we found that populations of an ecologically and commercially important fish with a ∼30-day pelagic larval stage are connected by dispersive currents from narrow windows of time. Identifying both the types and timing of currents that drive population connectivity can allow for better conservation and management decisions such as identifying the appropriate size and spacing of protected areas (Baetscher et al., 2019; Carr et al., 2017) or determining the boundaries of populations and management units (Waples & Gaggiotti, 2006). Ultimately, this framework, or a similar implementation, could allow for real-time predictions of recruitment (*e.g.*, integrating remote sensing and *in situ* observational data to forecast recruitment), which could greatly aid fisheries and conservation efforts. Scaling up these analyses to multiple species within a community could not only allow for greater generalizations, but could also identify the drivers, and their relative contributions, of population connectivity within and among aquatic systems.

## Acknowledgments

For sample collection, we thank the Michigan Department of Natural Resources, the Indiana Department of Natural Resources, the Wisconsin Department of Natural Resources, the Illinois Natural History Survey, and the Grand Traverse Band of Ottawa and Chippewa Indians for assistance. In particular, we thank Charles Roswell, Troy Zorn, Tammie Paoli, Dave Clapp, Pat O’Neil, Dave Fielder, David Schindelholz, Erik Olsen, Jayson Beugly, Josey Cline, Taylor Senegal, and Scott Koenigbauer. Sara Prendergast assisted with the Lagrangian particle dispersion model simulations. Field collections of yellow perch were authorized by issuance of state-specific collector’s permits and approved by Purdue University’s Animal Care and Use Committee protocol 1112000400. We thank Crow White for helpful discussions and Eric Crandall for reviewing an earlier draft of this manuscript. Any use of trade, product, or firm names is for descriptive purposes only and does not imply endorsement by the U.S. Government. This work was funded by the Great Lakes Fishery Commission (Project ID:2018_CHR_44072).

## Author Contributions

MRC and TOH designed the study, CS collected samples with agency assistance, AA and WL performed molecular work, CS, AA, and MRC analyzed the data, MDR performed the biophysical modeling, MRC designed and implemented the eco-genetic model, and CS and MRC wrote the manuscript with input from all authors.

## Data Availability Statement and Benefit Sharing

All RAD-Seq data will be made available at the NCBI Short Read Archive: Accession Numbers: SRXXXX-SRXXXX. All filtering, population structure analyses, and eco-genetic modeling code is hosted at GitHub: https://github.com/ChristieLab/yellow_perch_dispersal and all genetic, biophysical, and eco-genetic data will be posted on Dryad: D_XXXX.

## Notes

### Competing Interest Statement

The authors have declared no competing interest.

