## Supplemental Materials for "Dispersive currents from narrow windows of time explain patterns of population connectivity in an ecologically and economically important fish"

Disclaimer: Any use of trade, product, or firm names is for descriptive purposes only and does not imply endorsement by the U.S. Government

Claire Schraidt, Amanda S. Ackiss, Wesley A. Larson, Mark D. Rowe, Tomas O. Höök, Mark R. Christie

**Supplementary Methods and Materials:**

*Permutation test*

A permutation test was used to determine whether regional genetic diversity estimates (*H_o_* and *A_r_*) between Green Bay and the main basin differed from one another. Briefly, a test statistic was created as the mean of the 18 main basin genetic diversity estimates minus the mean of the 5 Green Bay genetic diversity estimates (Table 1; young-of-year samples omitted). Next all of the estimates were randomly assigned to one of two populations with samples sizes of 18 and 5 without replacement, the test statistic calculated, and the process repeated 100,000 times. Lastly, the empirical value was compared to the permuted distribution of test statistics to calculate a probability of the observed difference occurring by chance (Figure S2). This process was performed separately for two estimates of genetic diversity (observed heterozygosity and allelic richness).

*Biophysical models*

The FVCOM is an unstructured grid, finite-volume, free surface, three-dimensional primitive equation ocean model that solves the momentum, continuity, temperature, salinity, and density equations (Chen, Liu, and Beardsley 2003). The unstructured grid of FVCOM conforms to complex coastline morphologies and allows for increased grid resolution in regions of interest. Turbulence closure was implemented through the MY-2.5 scheme for vertical mixing (Galperin et al. 1988), and the Smagorinsky scheme for horizontal mixing (Smagorinsky 1963). FVCOM has been previously implemented for the Great Lakes yielding accurate predictions of temperature, water levels, and currents (Anderson et al. 2015; Anderson, Schwab, and Lang 2010; Anderson and Schwab 2013; Bai et al. 2013). Skill assessment of the model showed improved simulation of currents and surface temperature in comparison to the previous generation of Great Lakes operational forecast models, and it is currently used as NOAA’s next-generation Great Lakes Operational Forecast System. We define the epilimnion as the upper layer, or surface-mixed-layer, of a thermally stratified lake, which was an outcome of the vertical turbulent diffusivity simulated by FVCOM and the metalimnion as the layer below the epilimnion, separating the epilimnion from the hypolimnion, characterized by a strong temperature gradient and low turbulent diffusivity.

*Additional goodness of fit measures*

To provide an additional estimate the goodness of fit, we first standardized estimates of the slope to have a maximum value of 1 because many values were greater than 1 (*i.e.*, slope estimates of 0.8 and 1.2 were standardized to a value of 0.8 as they are equally distant from the 1:1 line and the direction of deviation does not matter for goodness of fit). We next added the standardized slope and *R^2^* values together assuming an additive relationship and that a perfect fit between the estimated and predictive value would have a slope of 1 and an *R^2^* of 1 (all points would fall on the 1:1 line). Thus, our goodness of fit values could range from < 0 (there were a few negative slopes) to 2 (perfect fit). For each combination of parameter values and release dates, we calculated the average predictive ability (goodness of fit averaged over 100 simulations) of the eco-genetic model at explaining the empirical values. We also examined the relationship between larval connectivity (measured as the number of grid cells with particles originating from cell *i*, averaged for all values of *i*), larval retention (measured as the number of particles that originated and remained in cell *i*, averaged for all values of *i*), and goodness of fit.

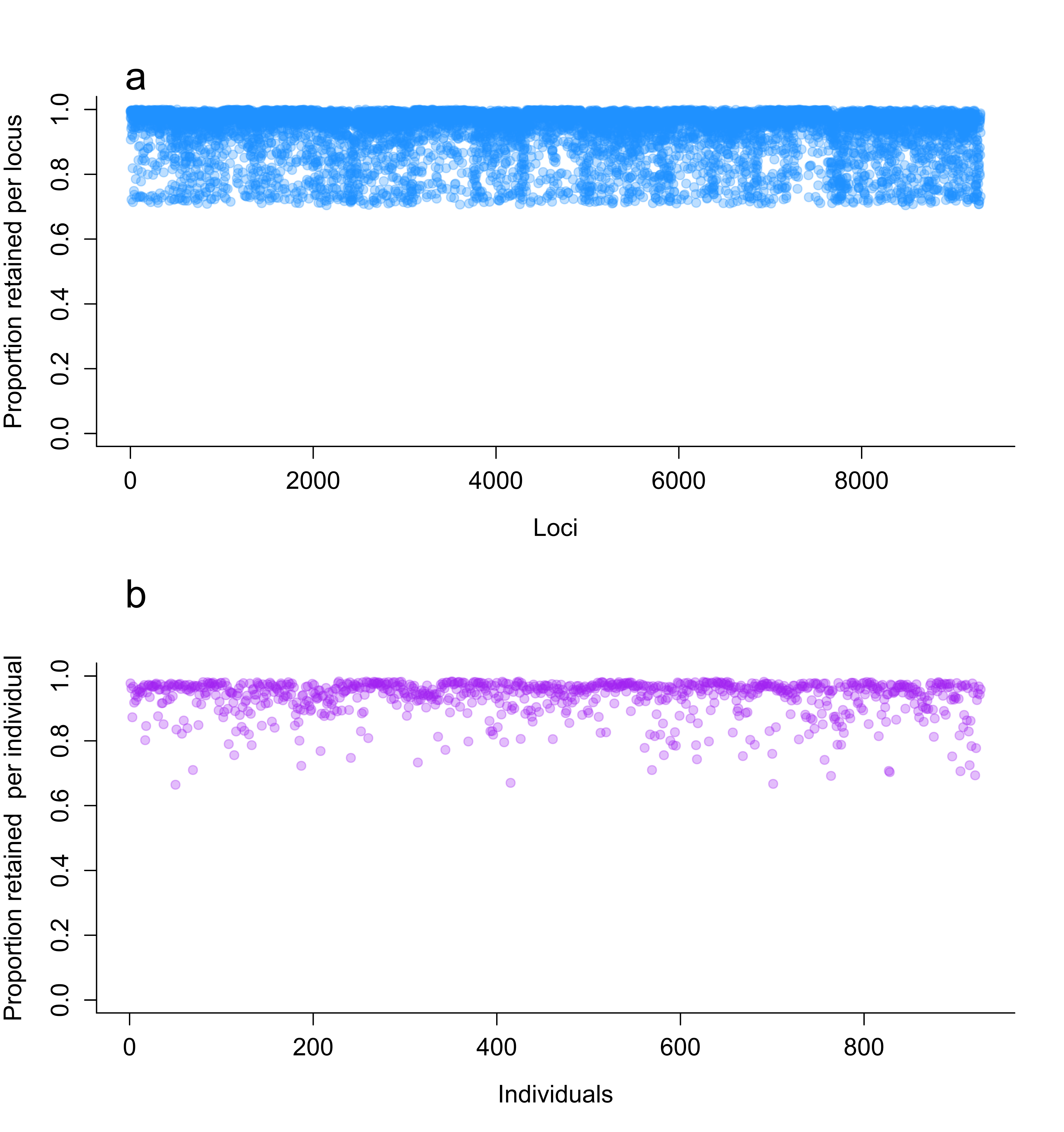

**Figure S1**: Percent of data retained (1 - %missing) per locus (a) and per individual (b). The average amount of missing data per locus was 6.48% across all individuals. The average amount of missing data per individual across all loci was also 6.48%. The average % missing values are identical due to the filtering options used but notice that the variation differs.

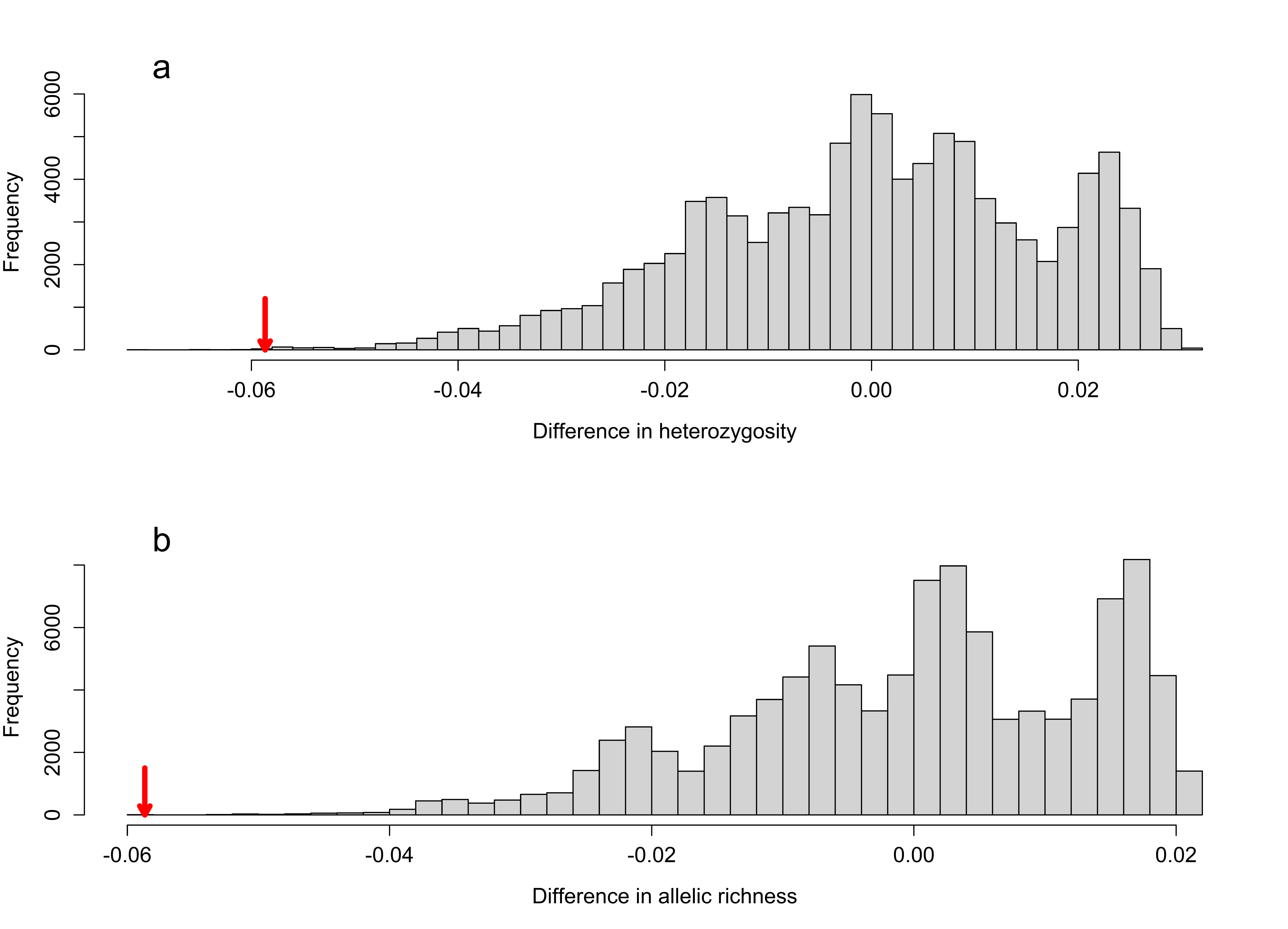

**Figure S2:** Results of permutation tests for determining whether observed heterozygosity (a) and allelic richness (b) was higher in main basin yellow perch populations than Green Bay yellow perch populations. Young of year fish were removed from this analysis. The mean difference in observed heterozygosity and allelic richness between main basin and green bay populations was -0.06 and -0.058, respectively (illustrated with red arrows). A total of 100,000 permutations was performed for each analysis. The p-value for observed heterozygosity and for allelic richness was < 0.001, however the observed difference is so small to be of little biological significance.

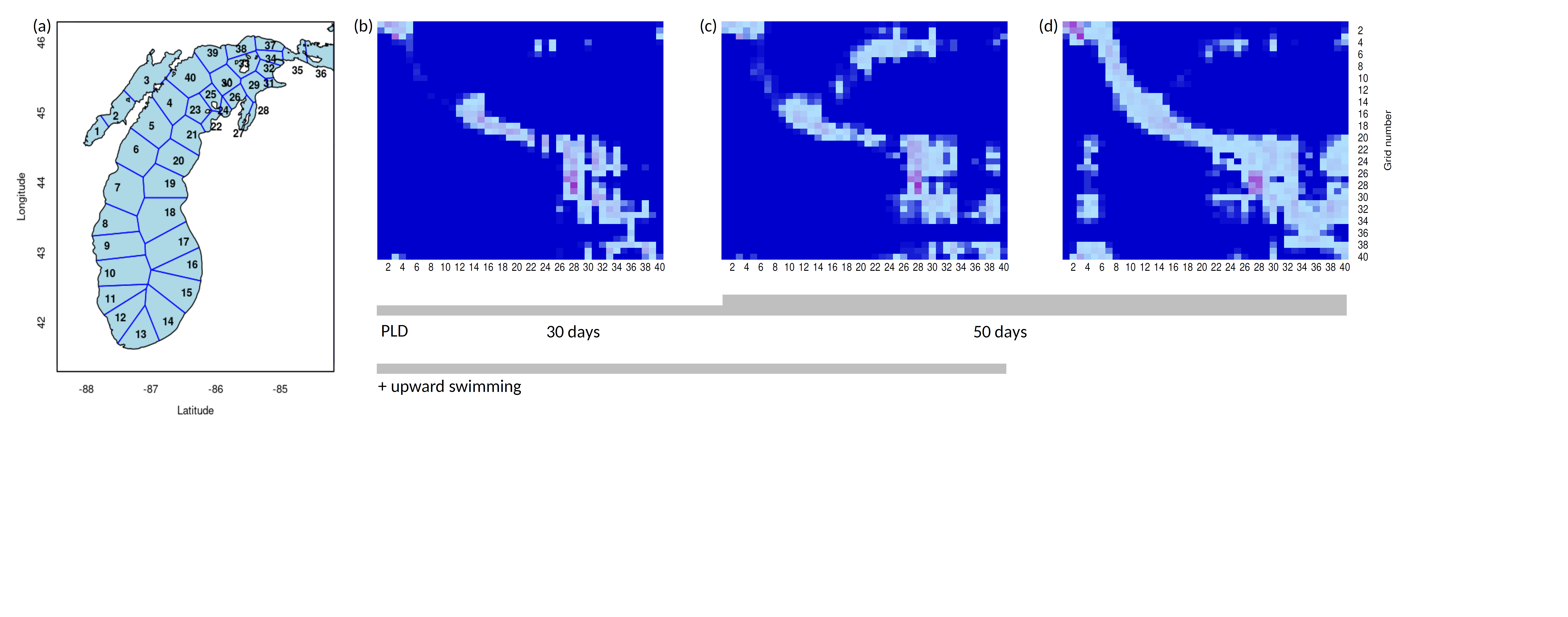

**Figure S3:** Current-driven patterns of Lake Michigan population connectivity as identified via biophysical models. (a) In order to track simulated particles, Lake Michigan was divided into 40 roughly equally sized polygons. Connectivity matrices from particles released June 30, 2019 illustrate population connectivity for particles with a simulated 30-day pelagic larval duration (PLD) and with upward swimming (0.0003 m/s) (b), particles with a simulated 50-day pelagic larval duration and upward swimming (c), and particles with a 50-day pelagic larval duration and no upward swimming (d). In these examples, connectivity increases with increases in the pelagic larval duration and neutral buoyancy (no upward swimming). Notice that there is 1.) high population connectivity within Green Bay, 2.) more population connectivity within northern than southern main basin sites, and 3.) fairly high larval retention and a general pattern of northward dispersal in the main basin.

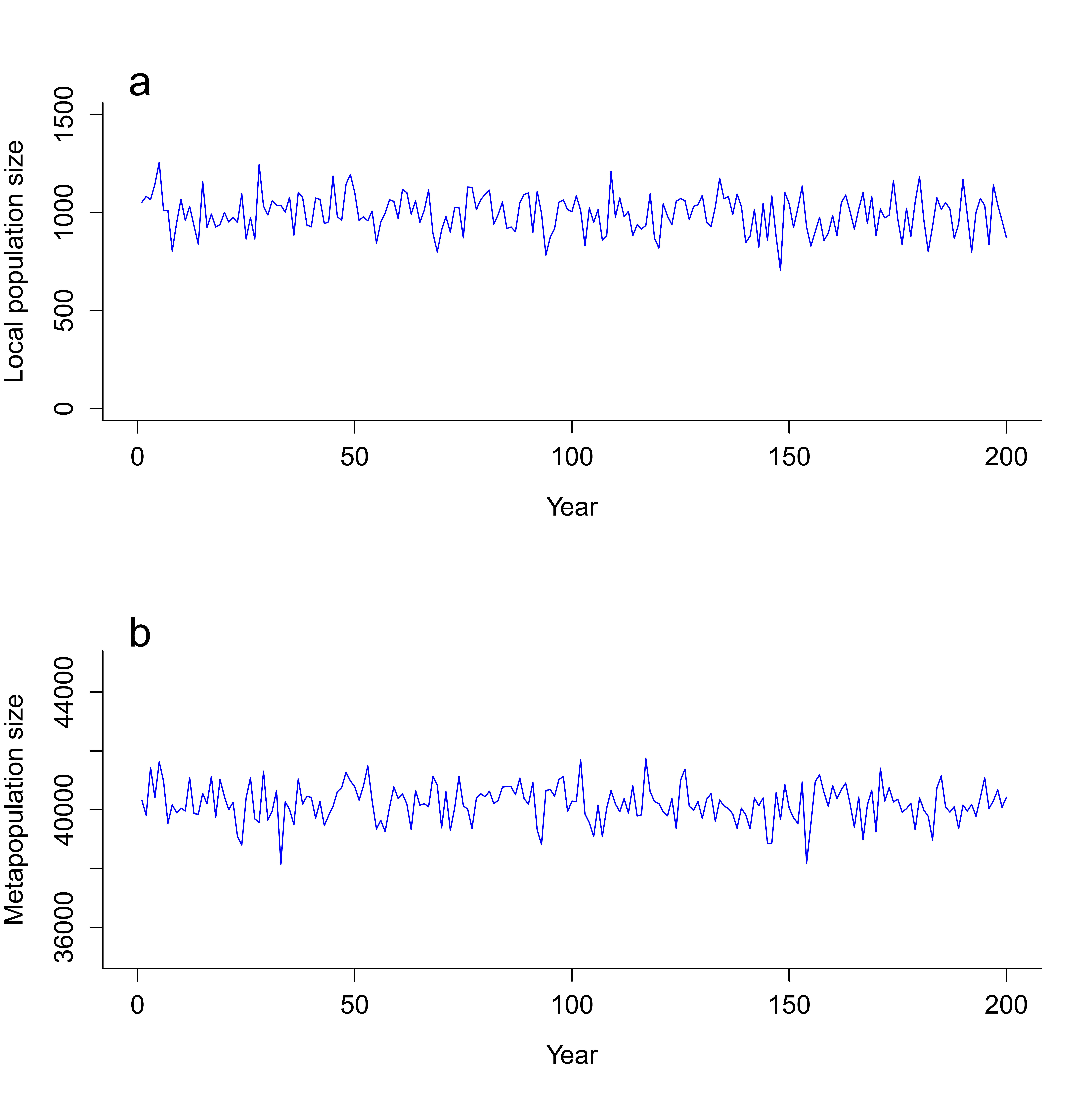

**Figure S4:** Examples of population dynamics through time in (a) one of 40 modeled local populations and (b) across all 40 local populations (*i.e*., the metapopulation). Notice that there was variation in each local population per year that, when summed over all local populations, resulted in fluctuations to the entire metapopulation.

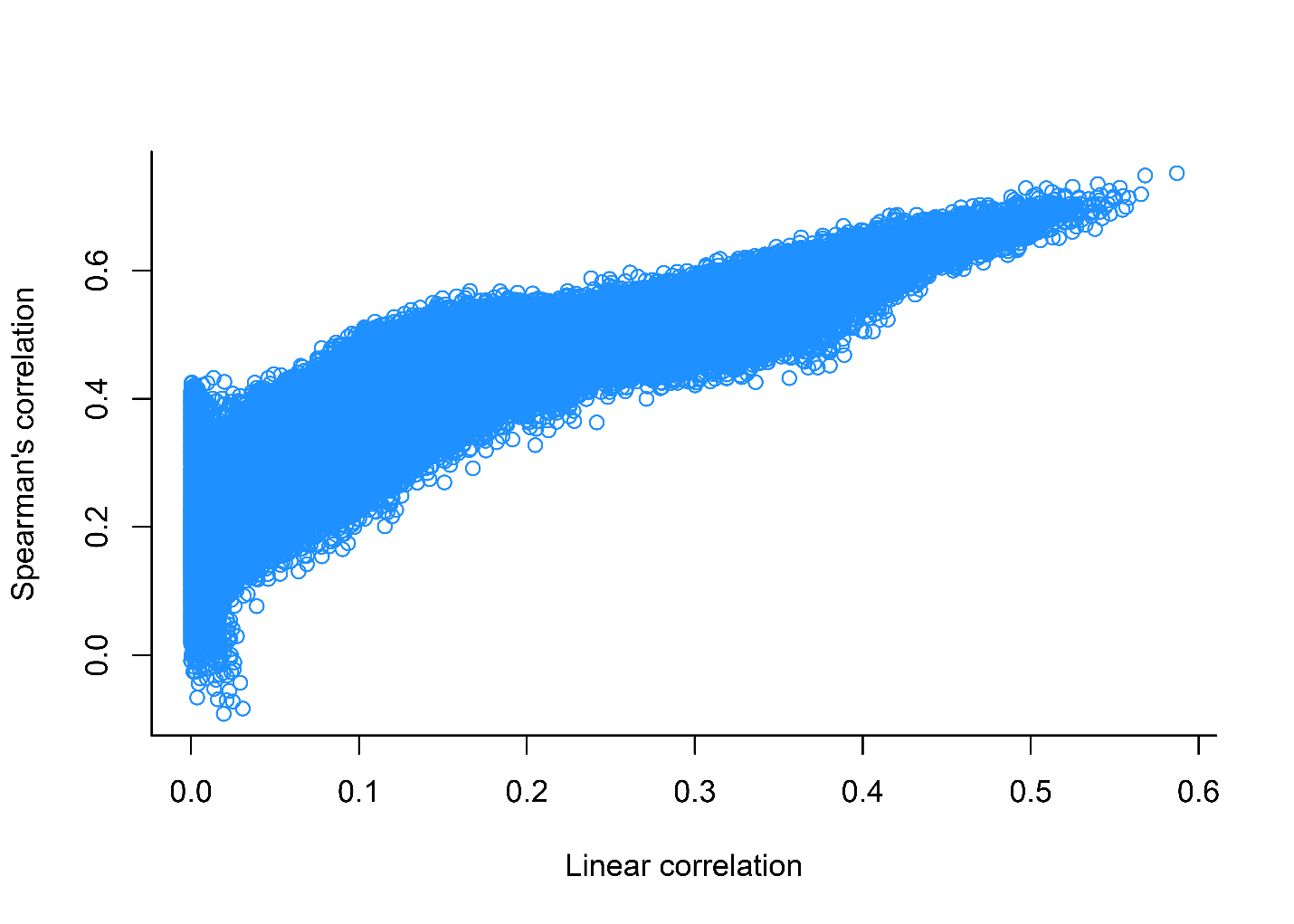
**Figure S5:** Relationship between linear correlation and Spearman’s rank correlation for estimating goodness of fit. The results were largely similar such that we relied on linear correlation estimates for the remainder of the analyses.

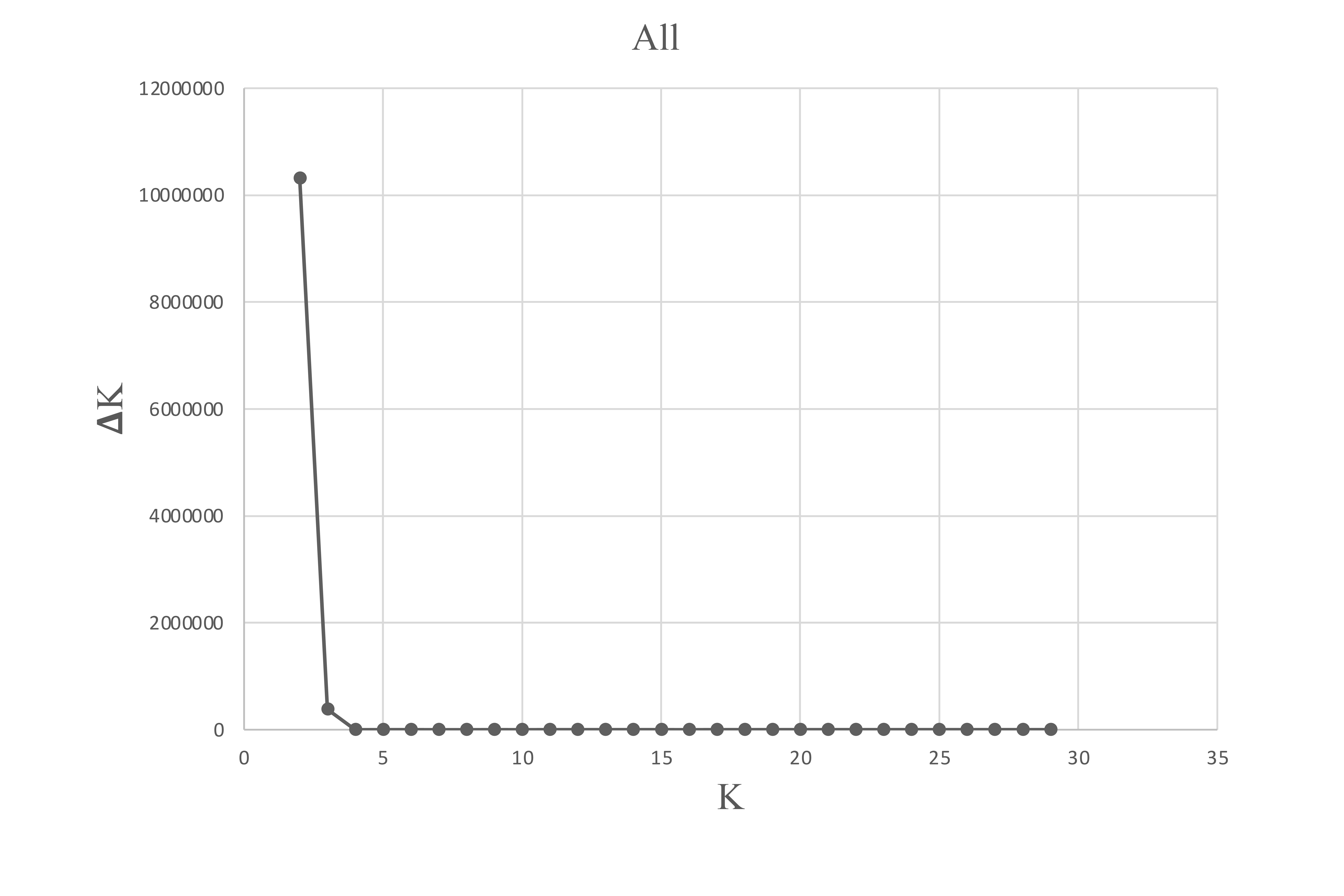

**Figure S6:** Changes in K values for all collection sites using STRUCTURE suggesting that no improvements were gained for larger numbers of clusters (see main text for details) and that it is sufficient to highlight the two main clusters illustrated in figure 1.

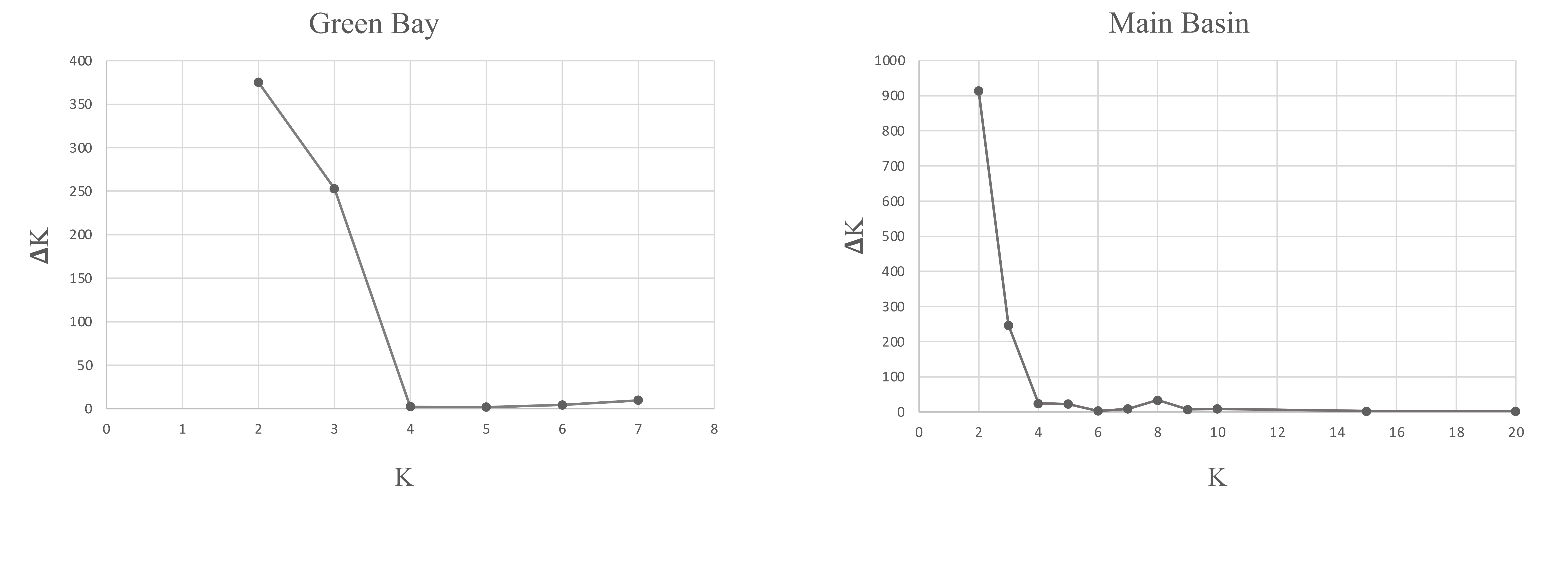

**Figure S7:** Changes in K values for all Green Bay (left panel) and all main basin (right panel) analyzed separately with STRUCTURE. In Green Bay, STRUCTURE analysis revealed similar ∆K values for K = 2 and K = 3, however K = 3 appears to be better supported by the results from the principal coordinate and principal component analyses (Figure 2a,b,c). These three groupings represent Little Bay de Noc, Big Bay de Noc, and Southern Green Bay, respectively. When the main basin samples were run separately to determine fine-scale population structure, analysis revealed ∆K maxima at K=2, but visualizing these clusters revealed only subtle population structure between the northern and southern sites (Figure 2d,e,f)

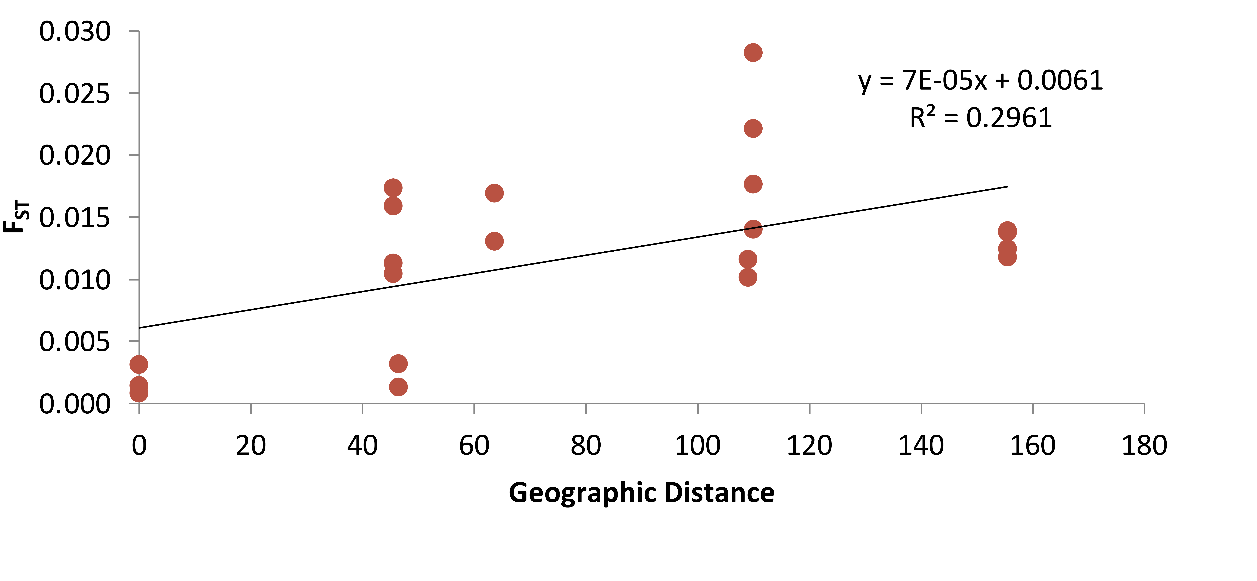

**Figure S8:** Isolation by Distance relationship for Green Bay collection sites. Plotted is the nearest along-shore distance (kilometers) versus pair-wise unbiased *F_ST_*. Mantel tests revealed that the slope is different than one (p = 0.05). Figure 2 also reveals differences among northern and southern Green Bay collection sites.

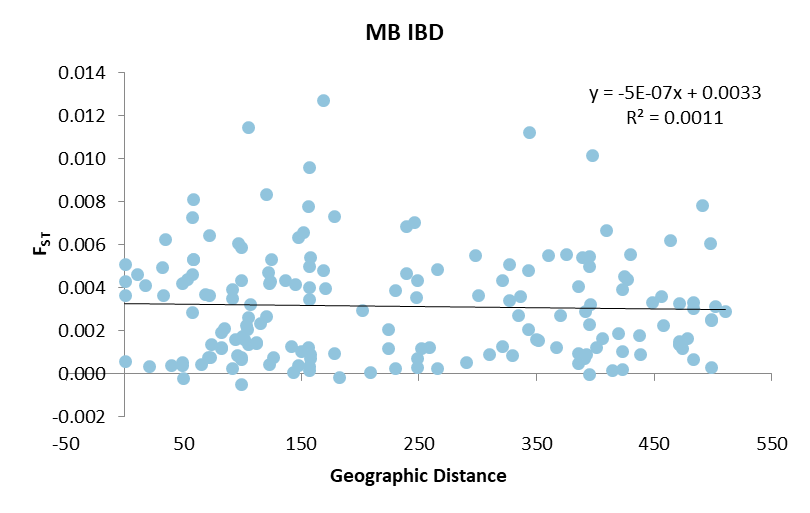

MB IBD

FST

| Geographic Distance | Y |
| --- | --- |
| 52.8109 | 0.0043763 |
| 32.0805 | 0.00366192 |
| 20.8852 | 0.000349882 |
| 17.5076 | 0.00414071 |
| 70.2738 | 0.0007827 |
| 49.5856 | -0.000186605 |
| 168.875 | 0.00483379 |
| 147.534 | 0.000419014 |
| 155.972 | 0.00122732 |
| 177.669 | 0.000979832 |
| 168.875 | 0.0127262 |
| 147.534 | 0.00635776 |
| 155.972 | 0.00780555 |
| 177.669 | 0.00734222 |
| 0 | 0.00363433 |
| 388.448 | 0.00540694 |
| 351.501 | 0.0015654 |
| 367.012 | 0.00124348 |
| 400.489 | 0.00123262 |
| 229.403 | 0.000244888 |
| 229.403 | 0.00389366 |
| 57.8807 | 0.00534149 |
| 98.5252 | 0.000782526 |
| 82.0275 | 0.00118746 |
| 48.3972 | 0.000388539 |
| 156.715 | 0.000151866 |
| 156.715 | 0.00344651 |
| 385.644 | 0.000476812 |
| 57.8807 | 0.00813333 |
| 98.5252 | 0.00433406 |
| 82.0275 | 0.00195542 |
| 48.3972 | 0.00420931 |
| 156.715 | 0.00404122 |
| 156.715 | 0.00962068 |
| 385.644 | 0.00406404 |
| 0 | 0.00510753 |
| 497.619 | 0.00607689 |
| 457.224 | 0.00227606 |
| 474.122 | 0.00120357 |
| 510.598 | 0.00289159 |
| 343.165 | 0.00209179 |
| 343.165 | 0.00483362 |
| 114.841 | 0.00237087 |
| 498.557 | 0.00250974 |
| 498.557 | 0.00031493 |
| 490.838 | 0.00783353 |
| 456.051 | 0.00360106 |
| 470.846 | 0.00326211 |
| 502.002 | 0.00313179 |
| 326.844 | 0.00343355 |
| 326.844 | 0.00510388 |
| 106.167 | 0.00322354 |
| 483.409 | 0.00306937 |
| 483.409 | 0.000699025 |
| 72.0852 | 0.00077722 |
| 426.621 | 0.00438453 |
| 390.171 | 0.00074196 |
| 405.536 | 0.00163503 |
| 438.427 | 0.000907958 |
| 265.833 | 0.000276565 |
| 265.833 | 0.0048579 |
| 38.8013 | 0.000399772 |
| 422.44 | 0.001052 |
| 422.44 | 0.0039529 |
| 84.1689 | 0.00213956 |
| 67.7514 | 0.00369975 |
| 57.8807 | 0.00534088 |
| 98.5252 | 0.000686087 |
| 82.0275 | 0.0012339 |
| 48.3972 | 0.000546583 |
| 156.715 | 0.000339274 |
| 156.715 | 0.00497961 |
| 385.644 | 0.000973709 |
| 0 | 0.000586093 |
| 0 | 0.00432765 |
| 498.557 | 0.00253891 |
| 483.409 | 0.00333475 |
| 422.44 | 0.000218041 |
| 124.375 | 0.00534328 |
| 73.2629 | 0.00137325 |
| 93.6875 | 0.00160646 |
| 141.299 | 0.00128443 |
| 122.877 | 0.000443923 |
| 122.877 | 0.00421301 |
| 289.996 | 0.000562676 |
| 157.519 | 0.000706943 |
| 157.519 | 0.00543161 |
| 390.888 | 0.00288788 |
| 395.947 | 0.00323252 |
| 328.772 | 0.000859702 |
| 157.519 | 0.000932344 |
| 95.8177 | 0.00608063 |
| 111.126 | 0.00140616 |
| 104.013 | 0.00208568 |
| 95.3211 | 0.000852621 |
| 104.994 | 0.00264579 |
| 104.994 | 0.0114406 |
| 334.089 | 0.00272217 |
| 57.342 | 0.00285824 |
| 57.342 | 0.00727826 |
| 448.106 | 0.00332371 |
| 429.427 | 0.00554275 |
| 370.019 | 0.00273039 |
| 57.342 | 0.00463803 |
| 144.695 | 0.00416519 |
| 245.895 | 0.00706195 |
| 208.414 | 7.86823e-05 |
| 223.846 | 0.00207291 |
| 258.571 | 0.00123799 |
| 98.6242 | -0.000490658 |
| 98.6242 | 0.00588628 |
| 143.18 | 7.47916e-05 |
| 248.087 | 0.000743202 |
| 248.087 | 0.0043368 |
| 252.072 | 0.00118524 |
| 247.777 | 0.00355431 |
| 181.782 | -0.000168514 |
| 248.087 | 0.000326901 |
| 150.065 | 0.00106306 |
| 201.371 | 0.00294581 |
| 119.994 | 0.00834803 |
| 121.518 | 0.00471981 |
| 120.107 | 0.00268084 |
| 123.147 | 0.0043265 |
| 71.8633 | 0.00363478 |
| 71.8633 | 0.00643882 |
| 300.151 | 0.00364248 |
| 90.6935 | 0.00392077 |
| 90.6935 | 0.000281286 |
| 414.395 | 0.000158749 |
| 394.992 | -2.1808e-05 |
| 335.845 | 0.00361642 |
| 90.6935 | 0.00350726 |
| 136.482 | 0.00434988 |
| 34.4701 | 0.00626134 |
| 169.754 | 0.00397228 |
| 397.159 | 0.0101675 |
| 359.692 | 0.00551148 |
| 375.41 | 0.00555491 |
| 409.346 | 0.0066542 |
| 238.859 | 0.00466094 |
| 238.859 | 0.00685345 |
| 9.92501 | 0.00464858 |
| 394.961 | 0.00500705 |
| 394.961 | 0.00231051 |
| 105.034 | 0.00138895 |
| 99.563 | 0.0017599 |
| 31.8179 | 0.0049451 |
| 394.961 | 0.00546088 |
| 297.447 | 0.00552201 |
| 343.622 | 0.0112303 |
| 151.578 | 0.00660439 |
| 309.735 | 0.000909263 |
| 463.391 | 0.0062179 |
| 419.306 | 0.00187335 |
| 437.533 | 0.00178375 |
| 477.614 | 0.00165734 |
| 320.44 | 0.00128237 |
| 320.44 | 0.00435015 |
| 111.559 | 0.00146557 |
| 471.113 | 0.00136114 |
| 471.113 | 0.00138751 |
| 64.5266 | 0.000441721 |
| 126.347 | 0.000761462 |
| 101.076 | 0.00162777 |
| 471.113 | 0.00150744 |
| 349.572 | 0.0016106 |
| 424.592 | 0.00455891 |
| 223.442 | 0.00119649 |
| 392.264 | 0.000897398 |
| 102.621 | 0.00225799 |

**Figure S9:** Isolation by Distance relationship for all main basin collection sites. Plotted is the nearest along-shore distance (kilometers) versus pair-wise unbiased *F_ST_*. Mantel tests revealed that the slope is not different than one (p = 0.36), suggesting that there is no relationship between along shore distance and genetic differentiation for main basin Lake Michigan yellow perch populations. Figure 2 reveals subtle differences among northern and southern main basin collection sites.

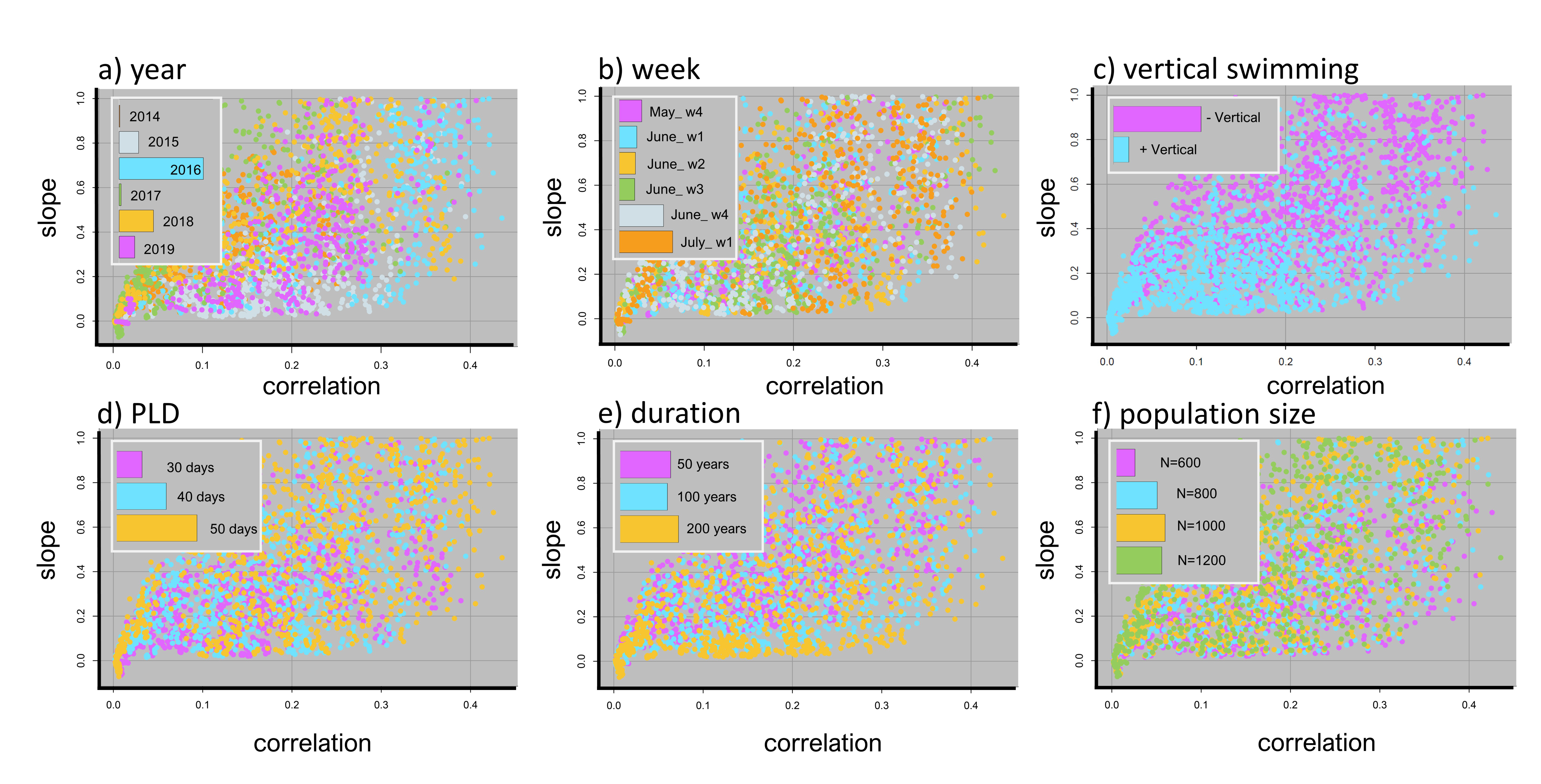

**Figure S10:** Drivers of population connectivity and genetic differentiation in main basin Lake Michigan yellow perch. Points represent the average correlation and slope values between predictive (generated from the integrated biophysical eco-genetic model) and empirical *F_ST_* for 100 simulations per unique set of parameter values (Table 3). A perfect fit would between predictive and empirical values would lie on the 1:1 line $(y=x)$ and would have a correlation and slope equal to one. Colors represent the effect of particular parameters (across all sets of parameter values). Insets illustrate the relative contributions of particular parameter values contributing to the top 20% of model predictions. Across all parameters, the specific year and week that particles were released in the biophysical model had high predictive ability (a, b), as did vertical swimming ability (c). Pelagic larval duration, the number of years that the eco-genetic model was run (“duration”), and the local population size had lower predictive ability (d, e, f).

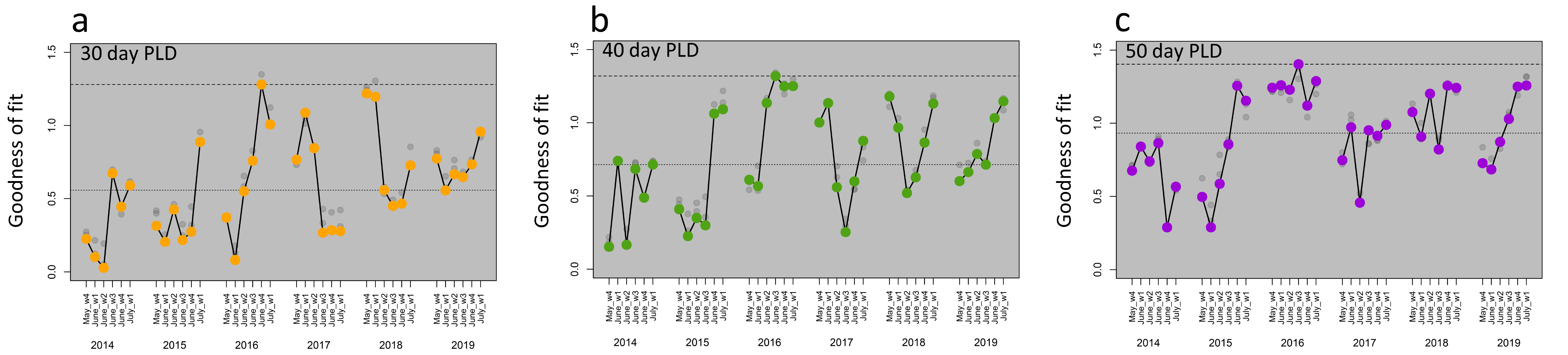

**Figure S11:** Drivers of population connectivity and genetic differentiation in Lake Michigan yellow perch. Points represent the average goodness-of-fit values between predictive and empirical *F_ST_* for 100 simulations per set of parameter values and are separated by particle release year and week. Colors represent pelagic larval durations of (a) 30 days (orange points), (b) 40 days (green points), and (c) 50 days (purple points) where particles were neutrally buoyant. The median and best goodness of fit values are illustrated with dashed, horizontal lines. For each combination of parameters, the eco-genetic model was run for 50, 100, or 200 years with the highest average goodness of fit indicated with the colored points (200 years in panels a-c) and the other two years indicated with smaller grey points. In some cases, the grey points lie directly under the colored point and are not visible. The specific release year and week explained the largest percentage of variation among all parameters examined. In this figure, goodness of fit was measured as described in Supplementary Methods.

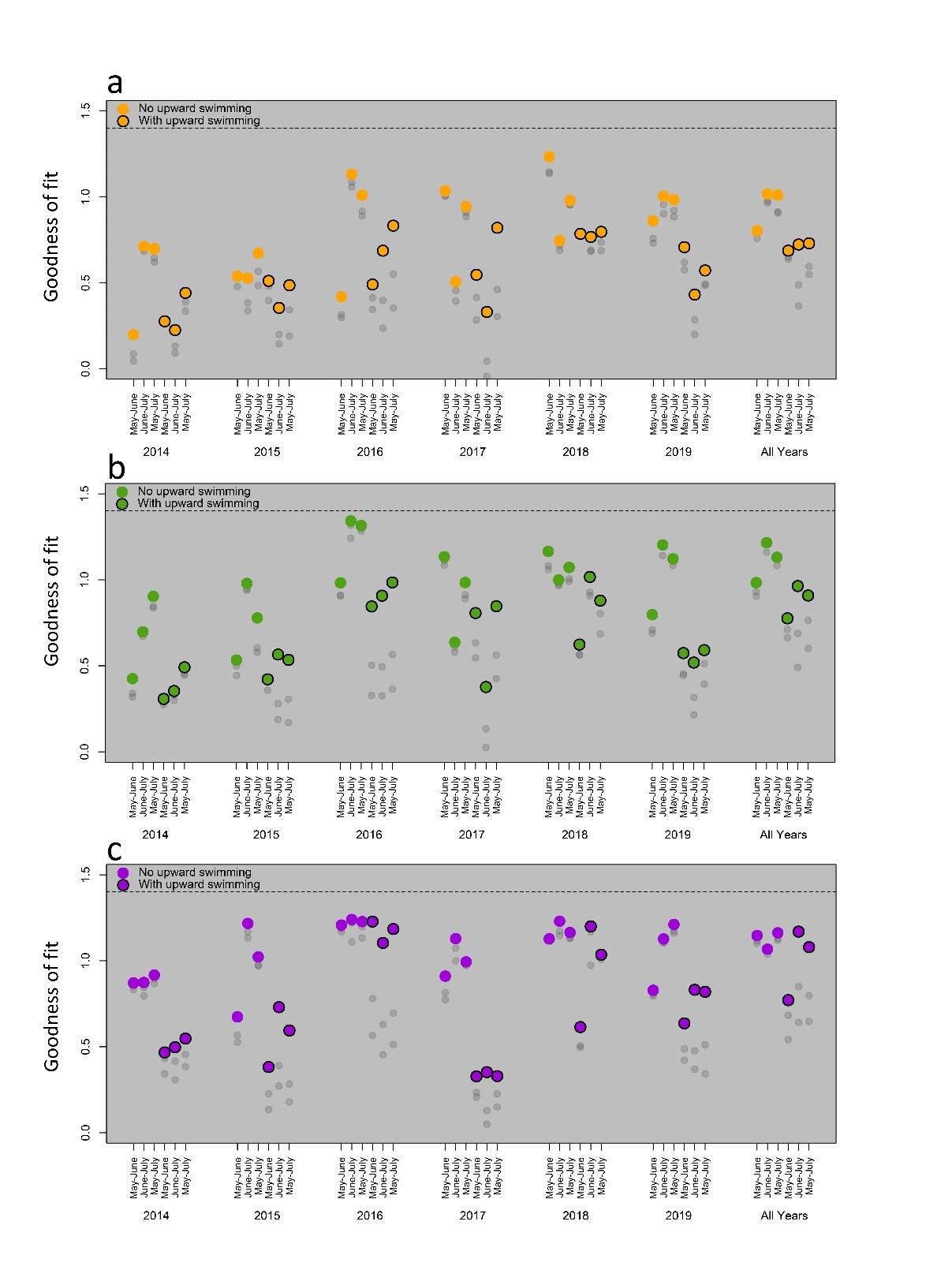

**Figure S12:** Goodness of fit between predictive *F_ST_* derived from our biophysical-ecogenetic model and empirical *F_ST_*. Here we evaluated whether varying connectivity matrices within a single eco-genetic simulation would improve predictive ability. We examined model runs where for each year of the biophysical model (2014-2019) where we: 1.) randomly selected a connectivity matrix from the first 3 release weeks for each year in the eco-genetic model ("May-June"), 2.) randomly selected a connectivity matrix for the last 3 release for each year in the eco-genetic model ("June-July"), and 3.) randomly selected a connectivity matrix from all 6 release weeks to be used for each year in the eco-genetic model ("May-June"). Lastly, we performed a similar analysis where for each combination of parameters and for each single release date, we randomly selected the year (2014-2019) from which drew the connectivity matrix ("All Years"). The horizontal dashed line represents the best goodness of fit from Figure 4.

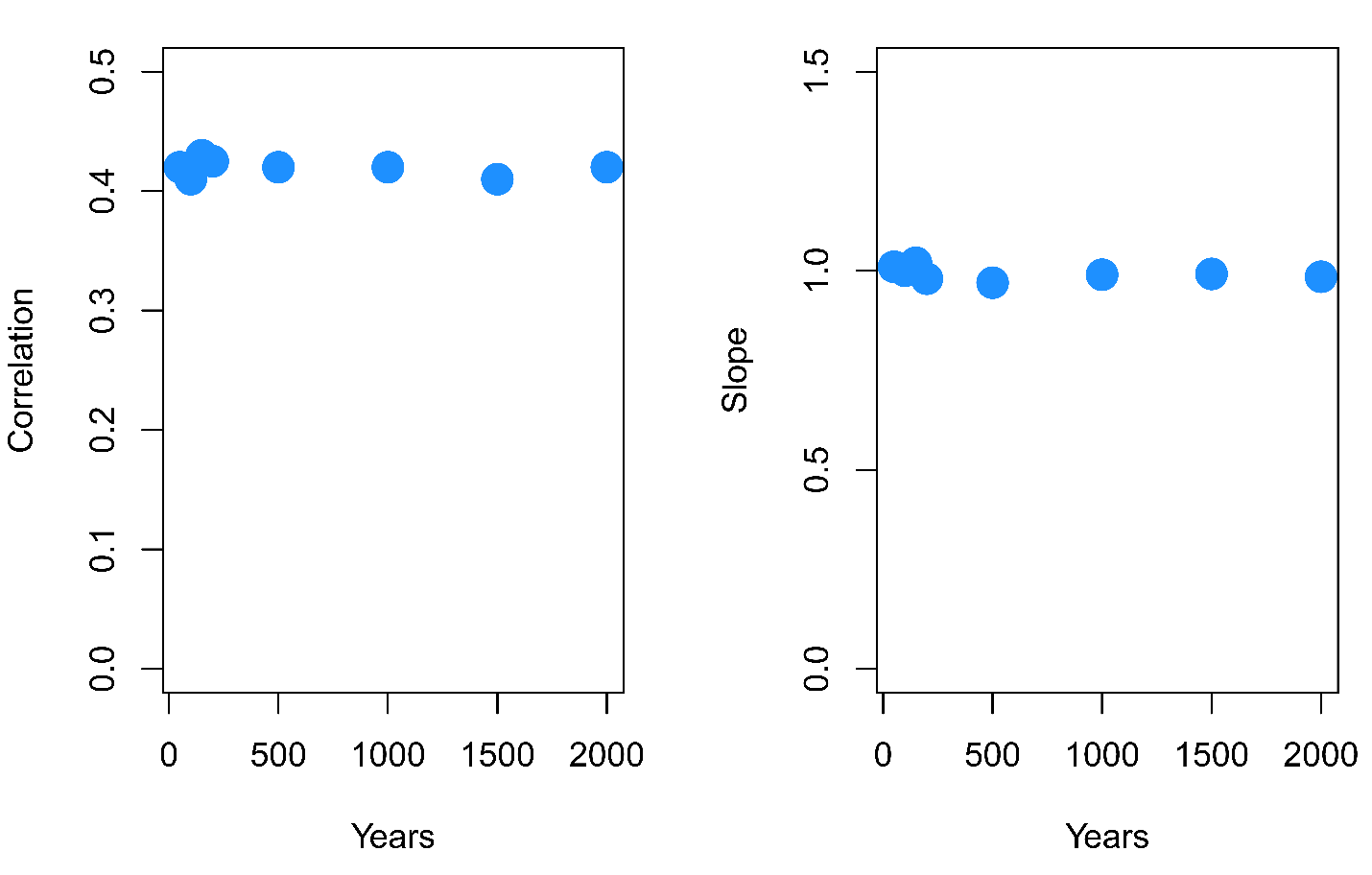
**Figure S13:** Examining the effect of long runs of the eco-genetic model on predictive ability. Here we picked one of the scenarios with highest predictive ability (week 5, 2016) and varied the number of years that the model was run, testing 50, 100, 150, 200, 500, 1000, and 2000 years). A total of 100 replicate simulations were run for each unique combination of parameters. Plotted are the mean correlation and slope for estimates of predictive estimates of *F_ST_* regressed against empirical estimates of *F_ST_*. Notice that varying the number of years that the eco-genetic model is run for has little effect on predictive ability (see also Figure 4 and Figure 5).

**Table S1:** Sampling details including collection site names, site ID, year collected ("Year"), Stage of individuals collected ("yoy = young of year"), Number of individuals collected ("Nc"), Number of individuals genotyped ("Ng"), Sex ratio (male:female), total length (range + SD), and weight (range + SD) for individuals measured.

| **Site** | **Site ID** | **Year** | **Stage** | **Nc** | **Ng** | **Sex Ratio** | **Total Length** | **Weight** |
| --- | --- | --- | --- | --- | --- | --- | --- | --- |
| Michigan City | MICYO | 2018 | yoy | 28 | 28 | na | 63 - 104 (7.9) | na |
| Michigan City | MIC19 | 2019 | adult | 50 | 46 | 21:29 | 158-365 (52.5) | 30-740 (180.8) |
| Michigan City | MIC18 | 2018 | adult | 100 | 50 | 22:28 | 128-332 (50.3) | 20-465 (111.3) |
| Saint Joseph | STJ18 | 2018 | adult | 130 | 50 | 45:05:00 | 121-296 (26.9) | 75-255 (45.5) |
| South Haven | SOH18 | 2018 | adult | 18 | 18 | na | 132-256 (33.5) | 20-170 (44.7) |
| South Chicago | SCH18 | 2018 | adult | 16 | 16 | na | 188-247 (18.7) | 62-173 (32.4) |
| Chicago | CHI18 | 2018 | adult | 13 | 13 | na | 145-312 (44.0) | na |
| North Chicago | NCH18 | 2018 | adult | 15 | 15 | na | 147-217(20.5) | na |
| Waukegan | WAK18 | 2018 | adult | 20 | 20 | na | 131-222 (27.4) | na |
| Grand Haven | GRH19 | 2019 | adult | 96 | 38 | na | na | na |
| Grand Haven | GRH18 | 2018 | adult | 109 | 50 | na | 143-225 (19.3) | 30-130 (24.6) |
| Ludington | LUD18 | 2018 | adult | 10 | 10 | 4:06 | 170-228 (16.8) | 20-95 (21.6) |
| Milwaukee | MIL19 | 2019 | adult | 32 | 32 | 15:17 | 166-357 (45.9) | na |
| Algoma | ALG18 | 2018 | adult | 1 | 1 | na | 194 (na) | 125 (na) |
| Suttons Bay | SUT18 | 2018 | adult | 32 | 32 | 4:25 | 190-319 (32.0) | 50-300 (70.36) |
| Northport | NPT18 | 2018 | adult | 34 | 34 | na | na | na |
| Charlevoix | CHX19 | 2019 | adult | 108 | 59 | na | na | na |
| Cheboygan | CHE18 | 2018 | adult | 41 | 40 | 7:34 | 212-269 (16.1) | 100-225 (37.4) |
| Naubinway | NUB18 | 2018 | adult | 72 | 72 | na | na | na |
| Manistique | MAN18 | 2018 | adult | 15 | 15 | na | na | na |
| Grand Haven | BDNYO | 2019 | yoy | 41 | 41 | na | 64-89 (5.7) | na |
| Grand Haven | BDN19 | 2019 | adult | 40 | 40 | na | 102-252 (35.5) | na |
| Little Bay de Noc | LBDYO | 2019 | yoy | 41 | 41 | na | 61 - 74 (3.4) | na |
| Little Bay de Noc | LBD19 | 2019 | adult | 40 | 40 | na | 104-315 (61.6) | na |
| Menominee | MEN19 | 2019 | adult | 49 | 49 | na | 102-264 (41.3) | na |
| South Green Bay | SGB19 | 2019 | adult | 84 | 50 | 34:16:00 | 134-311 (37.6) | na |
| South Green Bay | SGB18 | 2018 | adult | 82 | 60 | 28:32:00 | 100-305 (47.9) | na |

**Table S2:** Genetic diversity within sample sites and regions as measured by observed heterozygosity (*H_o_*), expected heterozygosity (*H_e_*), and allelic richness (*A_r_*).

| **Population** | **Region** | ***H_o_*** | ***H_e_*** | ***A_r_*** |
| --- | --- | --- | --- | --- |
| CHE18 | Main Basin | 0.229 | 0.235 | 1.238 |
| CHI18 | Main Basin | 0.216 | 0.218 | 1.229 |
| CHX19 | Main Basin | 0.234 | 0.231 | 1.234 |
| GRH18 | Main Basin | 0.222 | 0.227 | 1.229 |
| GRH19 | Main Basin | 0.293 | 0.236 | 1.239 |
| LUD18 | Main Basin | 0.222 | 0.212 | 1.228 |
| MAN18 | Main Basin | 0.224 | 0.226 | 1.235 |
| MIC18 | Main Basin | 0.225 | 0.226 | 1.229 |
| MIC19 | Main Basin | 0.226 | 0.226 | 1.228 |
| MICYO | Main Basin | 0.224 | 0.225 | 1.230 |
| MIL19 | Main Basin | 0.223 | 0.225 | 1.229 |
| NCH18 | Main Basin | 0.216 | 0.219 | 1.228 |
| NPT18 | Main Basin | NAN | 0.239 | 1.243 |
| NUB18 | Main Basin | 0.225 | 0.227 | 1.230 |
| SCH18 | Main Basin | 0.214 | 0.216 | 1.225 |
| SOH18 | Main Basin | 0.229 | 0.222 | 1.229 |
| STJ18 | Main Basin | 0.227 | 0.223 | 1.226 |
| SUT18 | Main Basin | 0.232 | 0.230 | 1.234 |
| WAK18 | Main Basin | 0.220 | 0.224 | 1.231 |
| BDN19 | Green Bay | 0.276 | 0.278 | 1.282 |
| BDNYO | Green Bay | 0.279 | 0.279 | 1.283 |
| LBD19 | Green Bay | 0.318 | 0.320 | 1.325 |
| LBDYO | Green Bay | 0.334 | 0.326 | 1.331 |
| MEN19 | Green Bay | 0.279 | 0.284 | 1.287 |
| SGB18 | Green Bay | 0.253 | 0.260 | 1.262 |
| SGB19 | Green Bay | 0.315 | 0.290 | 1.294 |

**Table S3:** Pair-wise estimates of *F_ST_* (Weir and Cockerham's unbiased estimator) for all 9,302 loci and all site comparisons. Numbers in parentheses represent the 95% confidence intervals) Full sample site information (including full site names) can be found in Tables 1 and S1 (Table S3 continued on next page).

|  | BDN19 | BDNYO | CHI18 | WAK18 |
| --- | --- | --- | --- | --- |
| BDN19 | 0 |  |  |  |
| BDNYO | 0.001, (-0.001,0.001) | 0 |  |  |
| CHI18 | 0.08, (0.077,0.082) | 0.081, (0.078,0.083) | 0 |  |
| WAK18 | 0.086, (0.083,0.09) | 0.087, (0.084,0.09) | 0.006, (0.004,0.007) | 0 |
| NCH18 | 0.076, (0.073,0.079) | 0.077, (0.074,0.08) | 0.004, (0.002,0.005) | 0.001, (-0.001,0.003) |
| SCH18 | 0.083, (0.08,0.087) | 0.085, (0.083,0.088) | 0.005, (0.003,0.007) | 0.001, (-0.002,0.002) |
| SGB18 | 0.013, (0.012,0.014) | 0.015, (0.014,0.015) | 0.101, (0.097,0.104) | 0.108, (0.105,0.112) |
| SGB19 | 0.015, (0.014,0.016) | 0.015, (0.014,0.016) | 0.106, (0.102,0.109) | 0.113, (0.11,0.116) |
| GRH18 | 0.093, (0.09,0.096) | 0.094, (0.09,0.097) | 0.006, (0.004,0.007) | 0.001, (-0.001,0.002) |
| GRH19 | 0.088, (0.085,0.091) | 0.089, (0.086,0.092) | 0.013, (0.012,0.015) | 0.007, (0.006,0.008) |
| MIC18 | 0.091, (0.088,0.094) | 0.092, (0.088,0.095) | 0.007, (0.005,0.008) | 0.002, (0.001,0.003) |
| MIC19 | 0.091, (0.087,0.094) | 0.092, (0.089,0.095) | 0.006, (0.004,0.007) | 0.001, (0.001,0.002) |
| LBD19 | 0.022, (0.021,0.023) | 0.023, (0.021,0.024) | 0.115, (0.112,0.118) | 0.123, (0.12,0.126) |
| LBDYO | 0.034, (0.032,0.035) | 0.031, (0.029,0.032) | 0.126, (0.122,0.129) | 0.134, (0.13,0.137) |
| MAN18 | 0.067, (0.064,0.069) | 0.068, (0.065,0.07) | 0.009, (0.007,0.011) | 0.006, (0.005,0.008) |
| CHE18 | 0.077, (0.075,0.08) | 0.078, (0.075,0.081) | 0.007, (0.006,0.009) | 0.004, (0.003,0.005) |
| CHX19 | 0.077, (0.074,0.079) | 0.077, (0.075,0.08) | 0.009, (0.008,0.011) | 0.005, (0.004,0.006) |
| MICYO | 0.088, (0.084,0.091) | 0.089, (0.086,0.092) | 0.005, (0.004,0.007) | 0.001, (0.001,0.002) |
| MEN19 | 0.012, (0.011,0.013) | 0.012, (0.011,0.013) | 0.1, (0.097,0.104) | 0.108, (0.104,0.112) |
| MIL19 | 0.086, (0.083,0.09) | 0.087, (0.084,0.09) | 0.006, (0.004,0.007) | 0.001, (-0.001,0.002) |
| STJ18 | 0.092, (0.088,0.095) | 0.092, (0.089,0.095) | 0.007, (0.006,0.008) | 0.002, (0.002,0.003) |
| LUD18 | 0.083, (0.079,0.086) | 0.084, (0.08,0.088) | 0.007, (0.005,0.009) | 0.002, (-0.001,0.004) |
| SOH18 | 0.086, (0.083,0.089) | 0.088, (0.084,0.091) | 0.007, (0.005,0.008) | -0.001, (-0.002,0.002) |
| SUT18 | 0.073, (0.071,0.076) | 0.075, (0.072,0.078) | 0.01, (0.008,0.011) | 0.006, (0.005,0.007) |
| NPT18 | 0.073, (0.071,0.075) | 0.074, (0.071,0.076) | 0.011, (0.01,0.013) | 0.007, (0.006,0.008) |
| NUB18 | 0.081, (0.078,0.084) | 0.083, (0.08,0.086) | 0.007, (0.006,0.009) | 0.002, (0.002,0.003) |

**Table S3:** Pair-wise estimates of *F_ST_* (Weir and Cockerham's unbiased estimator) for all 9,302 loci and all site comparisons. Numbers in parentheses represent the 95% confidence intervals) Full sample site information (including full site names) can be found in Tables 1 and S1 (Table S3 continued on next page).

|  | NCH18 | SCH18 | SGB18 | SGB19 | GRH18 |
| --- | --- | --- | --- | --- | --- |
| SCH18 | 0.001, (-0.001,0.003) | 0 |  |  |  |
| SGB18 | 0.095, (0.092,0.098) | 0.106, (0.103,0.111) | 0 |  |  |
| SGB19 | 0.101, (0.097,0.104) | 0.112, (0.108,0.115) | 0.006, (0.006,0.007) | 0 |  |
| GRH18 | 0.002, (0.001,0.003) | 0.001, (-0.001,0.002) | 0.113, (0.11,0.117) | 0.121, (0.118,0.125) | 0 |
| GRH19 | 0.008, (0.007,0.01) | 0.008, (0.007,0.009) | 0.109, (0.104,0.112) | 0.115, (0.111,0.118) | 0.004, (0.003,0.004) |
| MIC18 | 0.002, (0.001,0.003) | 0.002, (0.001,0.003) | 0.11, (0.107,0.114) | 0.119, (0.115,0.123) | 0.001, (-0.001,0.001) |
| MIC19 | 0.001, (-0.001,0.002) | 0.001, (-0.001,0.002) | 0.112, (0.108,0.115) | 0.119, (0.115,0.123) | -0.001, (-0.001,0.001) |
| LBD19 | 0.113, (0.11,0.115) | 0.119, (0.116,0.122) | 0.038, (0.036,0.039) | 0.018, (0.017,0.019) | 0.136, (0.133,0.14) |
| LBDYO | 0.124, (0.12,0.127) | 0.13, (0.126,0.134) | 0.048, (0.046,0.05) | 0.024, (0.023,0.025) | 0.148, (0.144,0.151) |
| MAN18 | 0.003, (0.001,0.004) | 0.005, (0.004,0.007) | 0.089, (0.085,0.092) | 0.094, (0.091,0.097) | 0.005, (0.004,0.006) |
| CHE18 | 0.002, (0.001,0.003) | 0.003, (0.002,0.004) | 0.098, (0.095,0.101) | 0.106, (0.102,0.108) | 0.003, (0.002,0.003) |
| CHX19 | 0.004, (0.003,0.005) | 0.004, (0.003,0.005) | 0.096, (0.093,0.099) | 0.106, (0.102,0.109) | 0.004, (0.004,0.004) |
| MICYO | 0.002, (0.001,0.003) | 0.001, (-0.001,0.002) | 0.11, (0.106,0.114) | 0.117, (0.113,0.12) | 0.001, (-0.001,0.001) |
| MEN19 | 0.096, (0.093,0.099) | 0.105, (0.102,0.109) | 0.006, (0.005,0.006) | 0.003, (0.002,0.003) | 0.117, (0.113,0.12) |
| MIL19 | 0.002, (0.001,0.003) | 0.001, (-0.002,0.002) | 0.108, (0.104,0.112) | 0.113, (0.11,0.117) | 0.001, (-0.001,0.001) |
| STJ18 | 0.002, (0.001,0.003) | 0.002, (0.001,0.003) | 0.112, (0.108,0.115) | 0.12, (0.117,0.123) | 0.001, (0.001,0.001) |
| LUD18 | 0.003, (0.001,0.005) | 0.003, (-0.001,0.005) | 0.107, (0.103,0.111) | 0.111, (0.108,0.115) | 0.003, (0.002,0.004) |
| SOH18 | 0.002, (0.001,0.003) | 0.001, (-0.001,0.003) | 0.108, (0.104,0.112) | 0.113, (0.11,0.117) | -0.001, (-0.002,-0.001) |
| SUT18 | 0.003, (0.002,0.004) | 0.005, (0.004,0.006) | 0.096, (0.092,0.099) | 0.103, (0.1,0.106) | 0.005, (0.004,0.005) |
| NPT18 | 0.006, (0.005,0.007) | 0.008, (0.006,0.009) | 0.094, (0.091,0.098) | 0.101, (0.098,0.104) | 0.006, (0.005,0.006) |
| NUB18 | 0.002, (0.001,0.003) | 0.002, (0.001,0.003) | 0.102, (0.099,0.106) | 0.111, (0.108,0.115) | 0.002, (0.001,0.002) |

**Table S3:** Pair-wise estimates of *F_ST_* (Weir and Cockerham's unbiased estimator) for all 9,302 loci and all site comparisons. Numbers in parentheses represent the 95% confidence intervals) Full sample site information (including full site names) can be found in Tables 1 and S1 (Table S3 continued on next page).

|  | GRH19 | MIC18 | MIC19 | LBD19 | LBDYO |
| --- | --- | --- | --- | --- | --- |
| GRH19 | 0 |  |  |  |  |
| MIC18 | 0.004, (0.004,0.005) | 0 |  |  |  |
| MIC19 | 0.004, (0.003,0.005) | 0.001, (-0.001,0.001) | 0 |  |  |
| LBD19 | 0.128, (0.125,0.131) | 0.134, (0.131,0.138) | 0.133, (0.13,0.137) | 0 |  |
| LBDYO | 0.14, (0.136,0.143) | 0.146, (0.142,0.15) | 0.145, (0.142,0.15) | 0.002, (0.002,0.003) | 0 |
| MAN18 | 0.011, (0.01,0.012) | 0.005, (0.004,0.006) | 0.006, (0.005,0.007) | 0.104, (0.101,0.106) | 0.115, (0.112,0.119) |
| CHE18 | 0.005, (0.005,0.006) | 0.003, (0.003,0.004) | 0.003, (0.003,0.004) | 0.119, (0.116,0.122) | 0.131, (0.128,0.134) |
| CHX19 | 0.006, (0.005,0.007) | 0.004, (0.004,0.005) | 0.004, (0.003,0.004) | 0.122, (0.118,0.125) | 0.134, (0.131,0.138) |
| MICYO | 0.005, (0.005,0.006) | 0.001, (0.001,0.002) | 0.002, (0.001,0.002) | 0.128, (0.125,0.132) | 0.14, (0.136,0.143) |
| MEN19 | 0.111, (0.107,0.114) | 0.114, (0.111,0.118) | 0.114, (0.111,0.118) | 0.021, (0.02,0.022) | 0.028, (0.026,0.028) |
| MIL19 | 0.005, (0.005,0.006) | 0.001, (0.001,0.002) | 0.001, (-0.001,0.002) | 0.126, (0.122,0.129) | 0.137, (0.133,0.141) |
| STJ18 | 0.005, (0.004,0.005) | 0.001, (0.001,0.002) | 0.001, (0.001,0.002) | 0.135, (0.132,0.139) | 0.147, (0.144,0.151) |
| LUD18 | 0.012, (0.01,0.013) | 0.003, (0.001,0.005) | 0.003, (0.002,0.004) | 0.115, (0.111,0.118) | 0.126, (0.122,0.129) |
| SOH18 | 0.006, (0.005,0.007) | 0.001, (-0.001,0.002) | 0.001, (-0.001,0.002) | 0.123, (0.12,0.126) | 0.134, (0.131,0.137) |
| SUT18 | 0.007, (0.006,0.008) | 0.005, (0.004,0.005) | 0.005, (0.004,0.006) | 0.116, (0.112,0.119) | 0.128, (0.124,0.131) |
| NPT18 | 0.008, (0.007,0.008) | 0.006, (0.005,0.007) | 0.006, (0.005,0.007) | 0.115, (0.112,0.118) | 0.127, (0.123,0.13) |
| NUB18 | 0.005, (0.004,0.005) | 0.002, (0.002,0.002) | 0.002, (0.001,0.002) | 0.127, (0.123,0.13) | 0.139, (0.135,0.143) |

**Table S3:** Pair-wise estimates of *F_ST_* (Weir and Cockerham's unbiased estimator) for all 9,302 loci and all site comparisons. Numbers in parentheses represent the 95% confidence intervals) Full sample site information (including full site names) can be found in Tables 1 and S1 (Table S3 continued on next page).

|  | MAN18 | CHE18 | CHX19 | MICYO | MEN19 |
| --- | --- | --- | --- | --- | --- |
| MAN18 | 0 |  |  |  |  |
| CHE18 | -0.001, (-0.002,0.001) | 0 |  |  |  |
| CHX19 | 0.001, (-0.001,0.002) | 0.001, (0.001,0.002) | 0 |  |  |
| MICYO | 0.005, (0.004,0.006) | 0.003, (0.002,0.004) | 0.005, (0.004,0.006) | 0 |  |
| MEN19 | 0.089, (0.085,0.092) | 0.101, (0.098,0.104) | 0.101, (0.098,0.104) | 0.112, (0.108,0.115) | 0 |
| MIL19 | 0.005, (0.004,0.006) | 0.003, (0.002,0.004) | 0.004, (0.003,0.005) | 0.001, (-0.001,0.001) | 0.109, (0.105,0.113) |
| STJ18 | 0.007, (0.006,0.008) | 0.004, (0.003,0.004) | 0.004, (0.004,0.005) | 0.001, (0.001,0.002) | 0.115, (0.111,0.118) |
| LUD18 | 0.008, (0.006,0.01) | 0.005, (0.003,0.006) | 0.006, (0.005,0.007) | 0.004, (0.002,0.005) | 0.105, (0.101,0.108) |
| SOH18 | 0.005, (0.004,0.007) | 0.002, (0.001,0.003) | 0.004, (0.004,0.005) | -0.001, (-0.001,0.001) | 0.108, (0.105,0.111) |
| SUT18 | 0.001, (-0.001,0.002) | 0.001, (-0.001,0.002) | -0.001, (-0.001,0.001) | 0.005, (0.004,0.006) | 0.098, (0.095,0.101) |
| NPT18 | 0.003, (0.002,0.004) | 0.002, (0.002,0.003) | 0.002, (0.002,0.002) | 0.006, (0.005,0.007) | 0.097, (0.094,0.1) |
| NUB18 | 0.002, (0.001,0.003) | 0.001, (-0.001,0.001) | 0.001, (0.001,0.002) | 0.002, (0.002,0.003) | 0.106, (0.103,0.11) |

**Table S3:** Pair-wise estimates of *F_ST_* (Weir and Cockerham's unbiased estimator) for all 9,302 loci and all site comparisons. Numbers in parentheses represent the 95% confidence intervals) Full sample site information (including full site names) can be found in Tables 1 and S1.

|  | MIL19 | STJ18 | LUD18 | SOH18 | SUT18 | NPT18 | NUB18 |
| --- | --- | --- | --- | --- | --- | --- | --- |
| MIL19 | 0 |  |  |  |  |  |  |
| STJ18 | 0.002, (0.001,0.002) | 0 |  |  |  |  |  |
| LUD18 | 0.005, (0.003,0.006) | 0.005, (0.004,0.007) | 0 |  |  |  |  |
| SOH18 | -0.001, (-0.001,0.001) | 0.001, (0.001,0.002) | 0.003, (0.001,0.005) | 0 |  |  |  |
| SUT18 | 0.004, (0.004,0.005) | 0.006, (0.005,0.006) | 0.007, (0.005,0.009) | 0.005, (0.004,0.006) | 0 |  |  |
| NPT18 | 0.006, (0.006,0.007) | 0.007, (0.006,0.007) | 0.012, (0.01,0.014) | 0.007, (0.007,0.008) | 0.001, (-0.001,0.002) | 0 |  |
| NUB18 | 0.002, (0.001,0.003) | 0.002, (0.002,0.003) | 0.005, (0.003,0.007) | 0.001, (0.001,0.002) | 0.001, (0.001,0.002) | 0.003, (0.002,0.003) | 0 |
